# Topoisomerase VI participates in an insulator-like function that prevents H3K9me2 spreading into euchromatic islands

**DOI:** 10.1101/829416

**Authors:** Louis-Valentin Méteignier, Cécile Lecampion, Florent Velay, Cécile Vriet, Laura Dimnet, Michel Térèse, Martin Rougée, Christian Breuer, Ludivine Soubigou-Taconnat, Keiko Sugimoto, Fredy Barneche, Christophe Laloi

## Abstract

The organization of the genome into transcriptionally active and inactive chromatin domains requires well-delineated chromatin boundaries and insulator functions in order to maintain the identity of adjacent genomic loci with antagonistic chromatin marks and functionality. In plants that lack known chromatin insulators, the mechanisms that prevent heterochromatin spreading into euchromatin remain to be identified. Here, we show that DNA Topoisomerase VI participates in a chromatin boundary function that safeguards the expression of genes in euchromatin islands within silenced heterochromatin regions. While some transposable elements are reactivated in mutants of the Topoisomerase VI complex, genes insulated in euchromatin islands within heterochromatic regions of the *Arabidopsis thaliana* genome are specifically downregulated. H3K9me2 levels consistently increase at euchromatin island loci and decrease at some TE loci. We further show that Topoisomerase VI physically interacts with S-adenosylmethionine (SAM) synthase MAT3, which is required for H3K9me2 deposition. Topoisomerase VI promotes MAT3 occupancy on heterochromatic elements and its exclusion from euchromatic islands, thereby providing a mechanistic insight into the essential role of Topoisomerase VI in the delimitation of chromatin domains.

## Introduction

The discovery of position effect variegation in *Drosophila melanogaster* paved the way towards revealing the importance of chromatin contexts in the regulation of gene expression (Wang *et al*, 2014; Muller, 1930). Since then, cytogenetic and molecular profiling of the epigenome, as well as topological analyses of chromatin architecture, have allowed the mechanisms involved in partitioning the gene-rich euchromatic fraction from the repeat-rich heterochromatic fraction to be elucidated. Large protein complexes specific to insulator DNA sequences contribute to partitioning chromatin domains with distinct identity at multiple scales. These complexes maintain the identity of adjacent genomic loci with antagonistic chromatin marks and functionality, and more globally influence the formation of long-range chromosomal interactions (Ali *et al*, 2016). Insulator binding proteins such as the CCCTC-binding factor CTCF, BEAF-32, CP190 and Mod have been best described in *Drosophila* where they play critical roles in the definition of chromatin and transcriptional status. In vertebrates, CTCF is the only known insulator binding protein homologue. CTCF is enriched at insulator DNA sequences that define large topological domains of the genome (Bickmore & van Steensel, 2013; Dekker & Misteli, 2015; Dixon *et al*, 2012) and, in some cases, define boundaries between adjacent chromatin domains with distinct features (Bonev & Cavalli, 2016). Surprisingly, CTCF orthologs cannot be identified in many eukaryotic organisms, including plants (Heger *et al*, 2012). Moreover, very few studies support the presence of insulator DNA sequences or insulator-like regions in plants (Wang *et al*, 2015a; Singer *et al*, 2011; Liu *et al*, 2017), and insulator binding factors remain to be identified. This contrasts with the observation that *Arabidopsis thaliana* and other plant species display highly indexed chromatin states along the genome, with well-defined chromatin signatures around transcriptionally active or repressed genes, as well as close relationships between chromatin composition and genome topology in the nuclear space (Liu & Weigel, 2015; Sequeira-Mendes & Gutierrez, 2016). In *Arabidopsis*, heterochromatin is found on hundreds of transposable elements (TEs) mostly confined within the pericentromeric regions and at a few knob structures that tend to associate through long-distance interactions in the nuclear space (Grob *et al*, 2013, 2014; Feng *et al*, 2014; Veluchamy *et al*, 2016; Liu *et al*, 2016). As a result, in *Arabidopsis* interphase nuclei most cytologically visible heterochromatin is condensed within 8 to 10 conspicuous foci that are referred to as chromocenters (Fransz & De Jong, 2011; Simon *et al*, 2015; Del Prete *et al*, 2014). Consistent with their heterochromatic nature, chromocenters are refractory to transcription and contain highly methylated DNA (Fransz *et al*, 2002) as well as histone modifications such as H3K9me2 and H3K27me2 (Soppe *et al*, 2002; Mathieu *et al*, 2005). Nonetheless, many expressed genes exhibiting euchromatic features appear to be located in close vicinity to large heterochromatic regions in the *Arabidopsis* genome, notably within the pericentromeric and knob regions (Lippman *et al*, 2004; Vergara & Gutierrez, 2017). The mechanisms by which gene-containing euchromatic islands (EIs) are insulated from neighboring heterochromatin regions and how their transcriptional capacities are preserved in such chromatin contexts are largely unknown. In this study, we have unveiled an essential function of the plant Topoisomerase VI complex in preserving the functional and structural identity of EIs.

DNA topoisomerases are enzymes that introduce transient DNA breaks to resolve topological constraints that arise during multiple cellular processes such as replication, transcription, recombination and chromatin remodeling. The plant Topo VI, a type II topoisomerase first identified in the archaeon *Sulfolobus shibatae* (Bergerat *et al*, 1994, 1997), was initially implicated in various biological processes involving endoreduplication, such as root hair growth (Schneider *et al*, 1998, 1997; Sugimoto-Shirasu *et al*, 2005), hypocotyl elongation (Sugimoto-Shirasu *et al*, 2005) and nodule differentiation (Yoon *et al*, 2014). Topo VI forms an A2B2 heterotetramer whose A and B subunits are encoded by single genes in *Arabidopsis*, *AtTOP6A*/*CAA39/AtSPO11-3*/*RHL2*/*BIN5*/*AT5G02820* and *AtTOP6B*/*RHL3/BIN3/HYP6/HLQ/AT3G20780*, respectively (Hartung & Puchta, 2001; Hartung *et al*, 2002; Yin *et al*, 2002; Mittal *et al*, 2014; Sugimoto-Shirasu *et al*, 2002). Two additional subunits named ROOT HAIRLESS 1 (RHL1/HYP7/AT1G48380) and BRASSINOSTEROID-INSENSITIVE 4 (BIN4/MID/AT5G24630) (Breuer *et al*, 2007; Kirik *et al*, 2007; Sugimoto-Shirasu *et al*, 2005) are essential for the *Arabidopsis* Topo VI function and appear to be evolutionarily conserved in plants and in other eukaryote groups, whilst their precise functions remain unclear. However, BIN4 shares sequence similarity with the C-terminal region of animal Topo IIα, which seems to have regulatory functions (Meczes *et al*, 2008; Onoda *et al*, 2014; Gilroy & Austin, 2011), and exhibits stable DNA binding *in vitro* (Breuer *et al*, 2007). Therefore, it has been proposed that BIN4 may have a regulatory role in the plant Topo VI complex, presumably by holding the substrate DNA through its AT-hook motif (Breuer *et al*, 2007; Kirik *et al*, 2007).

In recent years, evidence has accumulated that topoisomerases have more diverse and specialized functions than previously thought (Pommier *et al*, 2016). In particular, transcriptomic analyses of several Topo VI mutants revealed that Topo VI influences the expression of many nuclear genes, including genes regulated by phytohormones (Yin *et al*, 2002; Mittal *et al*, 2014) or by reactive oxygen species (Jain *et al*, 2006, 2008; Simkova *et al*, 2012). A function of *Arabidopsis* Topo VI as a chromatin-remodeling complex has also been speculated (Yin *et al*, 2002). This hypothesis has since been supported by the observation that loss of the Topo VI B subunit in *hlq* mutant plants leads to the mis-expression of numerous adjacent genes, hence possibly triggering positional or chromatin context dependent transcriptional defects (Mittal *et al*, 2014). This is further supported by the implication of the BIN4 subunit in heterochromatin organization, as observed by smaller and diffuse chromocenters in interphase nuclei of plants bearing the severe *mid* mutation (Kirik *et al*, 2007).

Here, we reveal that *Arabidopsis* Topo VI is required for chromocenter formation and for efficient silencing of some heterochromatic TEs, but has an antagonistic effect on genes localized in euchromatic islands (EIs). Downregulation of EI genes in Topo VI mutant plants is associated with an enrichment of the H3K9me2 heterochromatic mark. We further report that the BIN4 subunit of Topo VI directly interacts with S-adenosylmethionine (SAM) synthetase 3 / Methionine Adenosyl transferase 3 (MAT3). Similarly to Topo VI knockdown plants, *mat3* knockdown mutants exhibit de-repression of heterochromatic TEs and a decrease in H3K9me2. Furthermore, we show that MAT tethering to heterochromatin diminishes in a hypomorphic Topo VI mutant, leading to increased MATs localisation to EIs. This provides a mechanistic insight into the simultaneous local decrease of H3K9me2 in heterochromatin and increase of H3K9me2 at EIs in the Topo VI mutant. We therefore propose that Topo VI has a prominent role in the delimitation of chromatin boundaries, localizes SAM synthesis onto specific regions of the genome, and collectively has an essential role in the establishment of distinct chromatin domains.

## Results

### Topoisomerase VI is required for heterochromatin organization

Kirik *et al*. reported that interphase nuclei of the severe *mid* mutant in the BIN4/MID subunit presents smaller and less defined chromocenters (CCs) compared to the wild-type (wt), suggesting that heterochromatin organization is affected by the *mid* mutation (Kirik *et al*, 2007). However, this phenotype was not reported in the allelic *bin4-1* mutant, which also has a severe phenotype (Breuer *et al*, 2007). Therefore, to unequivocally confirm the role of the *Arabidopsis* Topo VI complex in nuclear organization, we analyzed the nuclear phenotypes of hypomorphic and amorphic mutants of the AtTOP6A subunit, *caa39* and *rhl2-1*, and of the BIN4/MID subunit, a *BIN4* knockdown line (*BIN4* KD, see below) and *bin4-1*, by DAPI DNA staining and immunolocalization of heterochromatin hallmarks. Both the *caa39* and *rhl2-1* mutants exhibited strong alterations in heterochromatin organization with largely decondensed chromocenters (Fig 1A, top panel, and Fig S1A). Likewise, nuclei of epidermal and mesophyll cotyledon cells from *BIN4* KD and *bin4-1* did not harbor conspicuous chromocenters (Fig S1A), as previously reported for the *mid* allelic mutants (Kirik *et al*, 2007). In contrast, the nuclear phenotype of shoot apical meristematic cells is similar in wt, *caa39*, *rhl2-1*, *bin4-1* and *BIN4* KD lines, with equal proportions of nuclei with conspicuous (type 1) or diffuse (type 2) chromocenter profiles (Fig S1A, meristem panel, and S1B). Consistent with its role in endocycles but not in mitosis (Breuer *et al*, 2007; Kirik *et al*, 2007; Sugimoto-Shirasu *et al*, 2002, 2005), these defects indicate that Topo VI is required for chromatin organization of differentiated cells, but less of actively dividing cells. Immunofluorescence analysis of the heterochromatin hallmark H3K9me2 confirmed the large extent of heterochromatin decondensation in *caa39* Topo VI mutant plants (Fig 1A). Immunoblot analyses further showed that the global level of H3K9me2 is not affected in *caa39* seedlings (Fig 1B). Likewise, 5-methylcytosine (5-meC) immunolabeling also revealed a dispersed signal in *caa39* nuclei (Fig 1C) whereas an anti-5-meC ELISA showed overall similar levels of 5-meC in wt and *caa39* as compared to *ddm1-8* seedlings (Fig 1D). These results suggest that the marked alteration of chromocenter morphology does not result from a global decrease in heterochromatin hallmarks in *caa39*.

**Figure 1.**
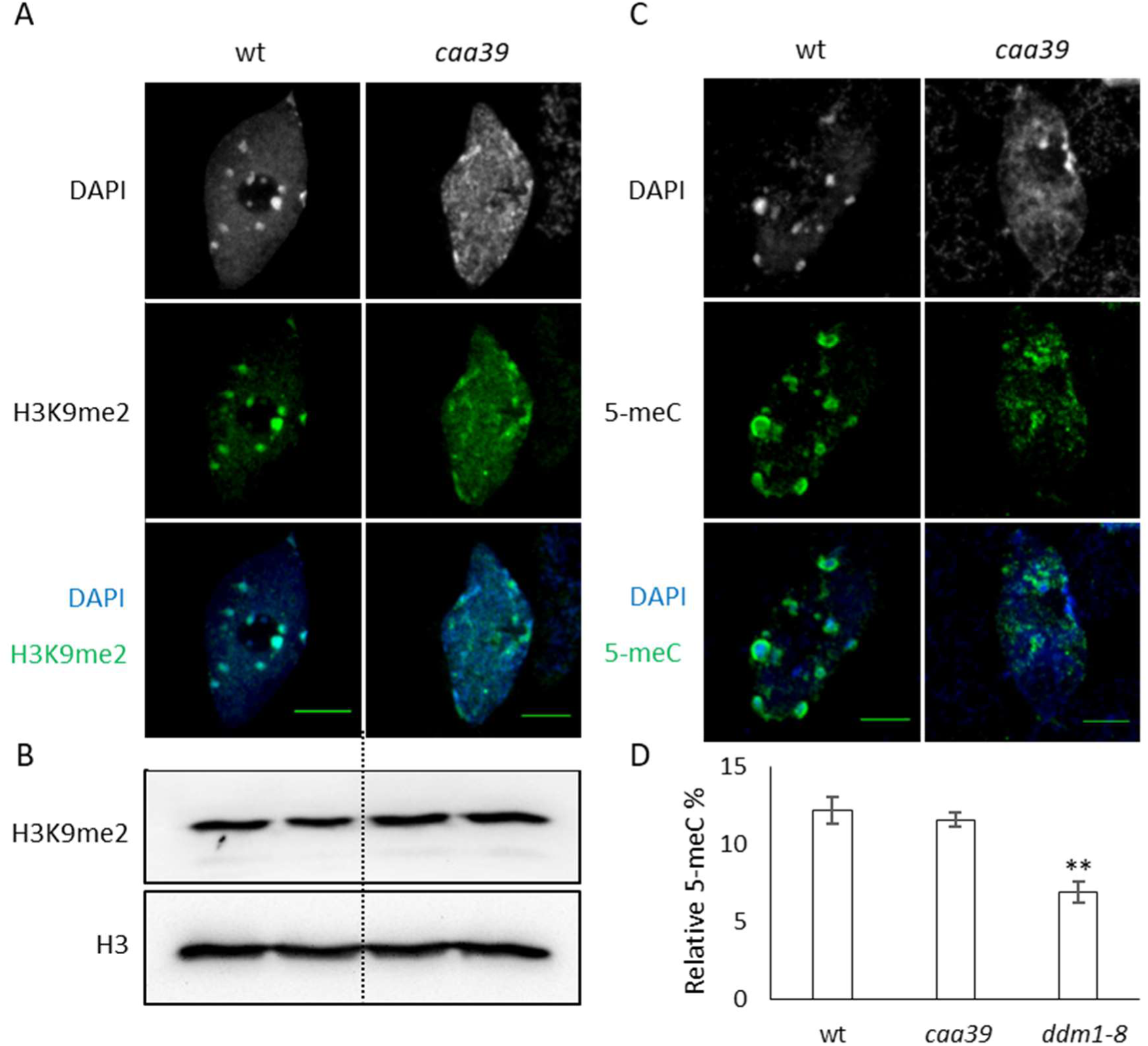
Topoisomerase VI is required for heterochromatin organization. (A) Representative nucleus (n>30) from 6 day-old wt and *caa39* cotyledon epidermal cells stained with DAPI and showing indirect immunolocalization of H3K9me2. Scale bar: 5 µm. (B) Two independently prepared nuclear extracts of wt and *caa39* were immunoblotted against H3 or H3K9me2, as indicated. (C) Same as (A) for 5-meC localization. (D) Elisa assay to quantify total 5-meC in 6 day-old cotyledons of wt, *caa39* and *ddm1-8*. **: *P* < 0.005 (Student’s *t*-test).

### Topo VI is required for the silencing of heterochromatic transposable elements

A role for *Arabidopsis* Topo VI in heterochromatin-dependent transcriptional gene silencing was highlighted by the reactivation of *TRANSCRIPTIONALLY SILENT INFORMATION* (*TSI*) in *mid* mutant plants (Kirik *et al*, 2007). However, reactivation was not observed in the *bin4-1* allelic mutant (Breuer *et al*, 2007). Therefore, to unambiguously assess the involvement of Topo VI in transcriptional silencing and get a more global understanding of Topo VI influence on TE repression, we performed a RNA-seq analysis of *caa39* and wt transcripts. Multiple heterochromatic TEs (176 TEs with log2(FC) > 2), particularly from the LTR/Gypsy, LTR/Copia and En-Spm/CACTA superfamilies (Underwood *et al*, 2017), are reactivated in *caa39* plants (Fig 2A, Appendix Table S1). Conversely, 91 TEs are repressed in *caa39* (log2(FC) < −2) and unlike reactivated TEs, these repressed TEs are rarely in the most inaccessible and repressive heterochromatin state 9 (Fig S2A). To test for TE silencing defects in other Topo VI mutants, we selected several de-repressed heterochromatic TEs loci (Appendix Table S1) for which robust primer design was feasible, and measured their relative transcript abundance by RT-qPCR in *rhl2-1* and *bin4-1* mutants along with the *caa39* and wt lines. A clear increase in TE transcript abundance was observed for all three tested Topo VI mutant lines (Fig 2B).

**Figure 2.**
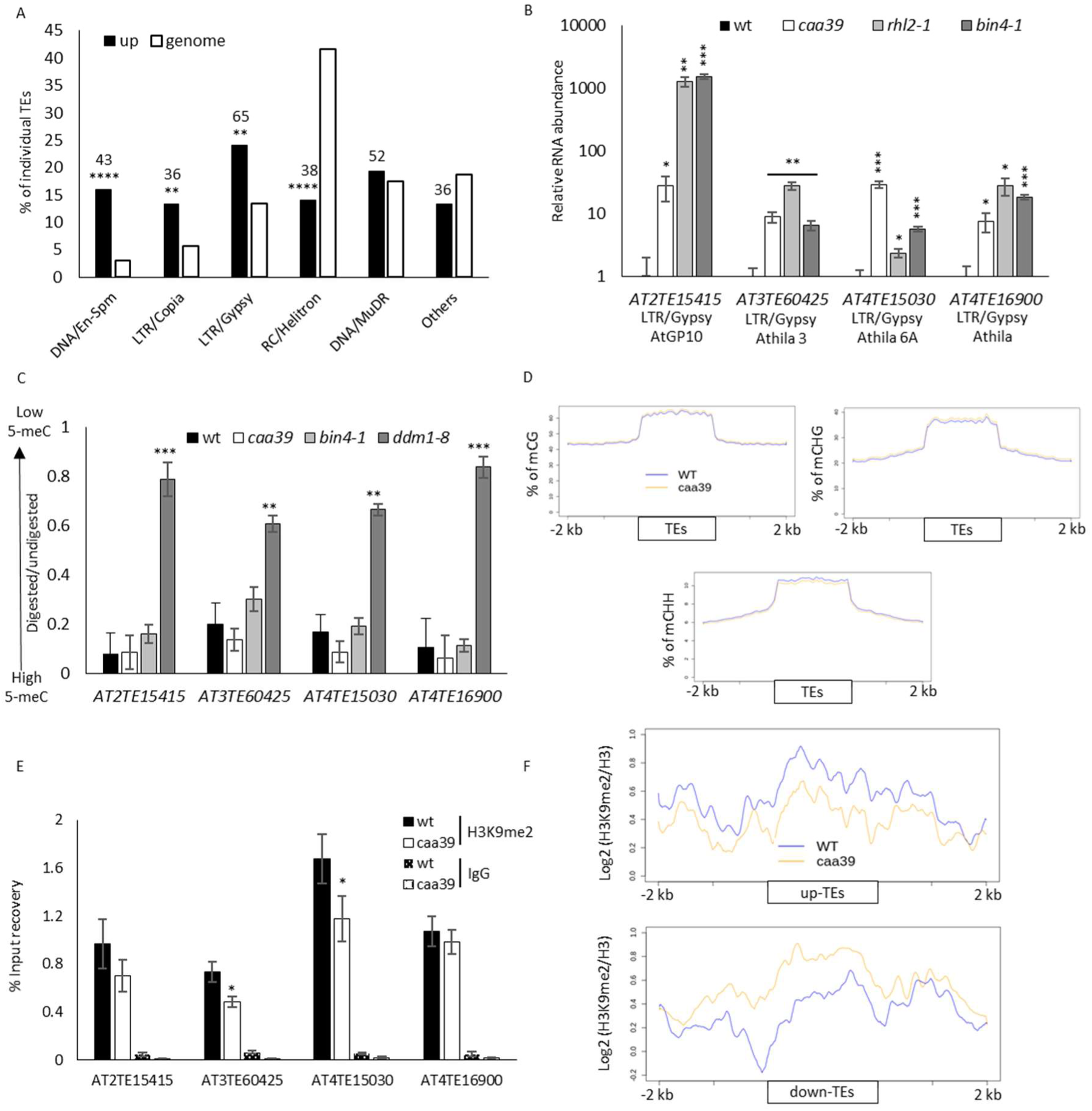
Topo VI is required for the silencing of transposable elements. (A) Bar chart showing the proportions of reactivated TE superfamilies in *caa39* compared to the general proportion of TEs in the genome. The relative percentage is shown for each superfamily and the absolute number of reactivated TEs is noted above each bar. **: P < 0.005; ****: P < 0.0001 (Chi-squared test). (B) RT-qPCR confirmation of the reactivation of selected TEs in *caa39*, *rhl2-1* and *bin4-1*. Error bars: ± SEM of three biological replicates. *: *P* < 0.05; **: *P* < 0.005; ***: *P* < 0.0005 (Student’s *t*-test). (C) DNA from indicated genotypes were extracted and digested with McrBC. The mean ratio of digested over undigested DNA from three biological replicates is shown. Error bars: ± SEM of three biological replicates. *: *P* < 0.05; **: *P* < 0.005; ***: *P* < 0.0005 (Student’s *t*-test). (D) Average distribution of methylated cytosine in CG, CHG and CHH contexts over up-or downregulated TEs. Two independent replicates for each genotype were performed. (E) ChIP-qPCR of H3K9me2 at selected TEs. Error bars: ± SEM of three biological replicates. *: *P* < 0.05; **: *P* < 0.005; ***: *P* < 0.0005 (Student’s *t*-test). (F) Average distribution of H3-normalized H3K9me2 along upregulated or downregulated TEs and 2 kb flanking regions. Two (H3) and three (H3K9me2) independent biological replicates for each genotype were performed.

Although we found no global decrease of H3K9me2 and 5-meC in *caa39* (Fig 1B and 1D), more subtle local changes could account for TE reactivation. We first assessed DNA methylation levels at individual TEs in Topo VI mutants as compared to wt and *ddm1-8* plants by digestion with the methylation-dependent restriction enzyme McrBC. As expected, very efficient digestion of TEs was observed in wt but not in *ddm1-8* plants, reflecting a nearly complete loss of DNA methylation over multiple TEs in this hypomethylated mutant line (Fig 2C). In sharp contrast, we observed wt levels of DNA methylation for all tested loci in *caa39* and *bin4-1* plants. This trend was confirmed in different sequence contexts (CG, CHG and CHH) by using the methylation-sensitive restriction enzymes HpaII, MspI and HaeIII (Fig S3). Similar DNA methylation levels of TEs in wt and *caa39* were then confirmed genome-wide by whole-genome bisulfite sequencing, in all three contexts (Fig 2D). We concluded that TE de-repression in Topo VI mutants could not be accounted for by a global decrease of DNA methylation in *cis*. Next, we measured the level of H3K9me2 at TEs, which could be performed only with the *caa39* hypomorphic mutant, owing to the extreme dwarf phenotype of the *rhl2-1* and *bin4-1* null mutants. ChIP-qPCR analyses revealed a modest decrease in H3K9me2 at some but not all tested TEs in *caa39* as compared to wt plants (Fig 2E). ChIP-seq analysis of H3K9me2 levels in wt and *caa39*, normalized to H3 levels in each line, confirmed the slight decrease of H3K9me2 at *AT3TE60425* and *AT4TE15030*, but not at *AT2TE15415* and *AT4TE16900* (Fig S2B-C), and revealed a consistent decrease or increase of H3K9me2 on upregulated or downregulated TEs, respectively (Fig 2F). Collectively, the results presented in figure 1 and 2 suggest that TE reactivation results from a combination of moderate and local decreases of H3K9me2 and heterochromatin reorganization. Moreover, the expression pattern of TEs correlates well with corresponding changes in their H3K9me2 content but not with DNA methylation changes.

### Unlike TEs, genes interspersed within pericentromeric and chromosome 4 knob large heterochromatin regions are massively downregulated in Topo VI mutants

We then used the online positional gene enrichment tool (De Preter *et al*, 2008) to investigate the genomic distribution of misregulated genes identified in our RNA-seq analysis of *caa39* (Appendix Table S2). This analysis revealed that the 500 most downregulated genes are strikingly over-represented in pericentromeric regions (PRs) and in the heterochromatic knob of chromosome 4 (*hk4S*, Fig 3A). In these regions, 94% (181/193) of the non-TE genes that are differentially expressed in *caa39* are downregulated (Appendix Table S2). In contrast, the 500 most highly upregulated genes displayed no preferential localization (Fig S4A). To determine whether this effect is robust in other Topo VI mutant lines, we first examined the expression profiles of *bin4-1* and a second allelic mutant, *bin4-2*, from microarray data that were generated during the initial characterization of these two allelic lines (Breuer *et al*, 2007). Despite the fact that *bin4-1* and *bin4-2* are knock-out mutants that have much more severe developmental defects than *caa39*, and although two different technical platforms have been used (RNA-seq for *caa39 vs* Affymetrix ATH1 microarrays for *bin4-1* and *bin4-2*), we found a good correlation between the different transcriptomes (Fig S4B). In particular, 91% (72/79) of the PR genes that are repressed in *caa39* and are detected in both RNA-seq and microarray experiments are also down-regulated in *bin4-1* or *bin4-2* (Fig 3B, Appendix Table S2). We examined further the expression of seven pericentromeric genes distributed over the five chromosomes and strongly repressed in *caa39*, by RT-qPCR in *caa39*, *bin4-1* as well as in *rhl2-1* plants. These genes were found to be downregulated in all mutants, except for *AT4G06634* and *AT4G07390* that were not significantly repressed in *rhl2-1* and *bin4-1* (Fig 3C). This could possibly be due to secondary effects of the *bin4-1* and *rhl2-1* amorphic mutations as compared to the less severe *caa39* mutation. In order to test this hypothesis, we took advantage of the availability of an *Arabidopsis BIN4* co-suppressed transgenic line identified during the process of generating lines overexpressing *BIN4-CFP*. Rather than exhibiting *BIN4* upregulation, this *BIN4* KD homozygous, monoinsertional transgenic line, shows downregulation of *BIN4* (Fig S5A-B) and develops a phenotype similar to *caa39* (Fig S5C). In this line, all tested pericentromeric genes were at least as much downregulated as in *caa39*, with a more pronounced effect than in the *bin4-1* and *rhl2-1* knockout mutants (Fig 3C). These observations indicate that Topoisomerase VI is required to maintain transcriptional control of both genes and TEs in pericentromeric and *hk4S* regions, possibly acting as a chromatin architectural factor.

**Figure 3.**
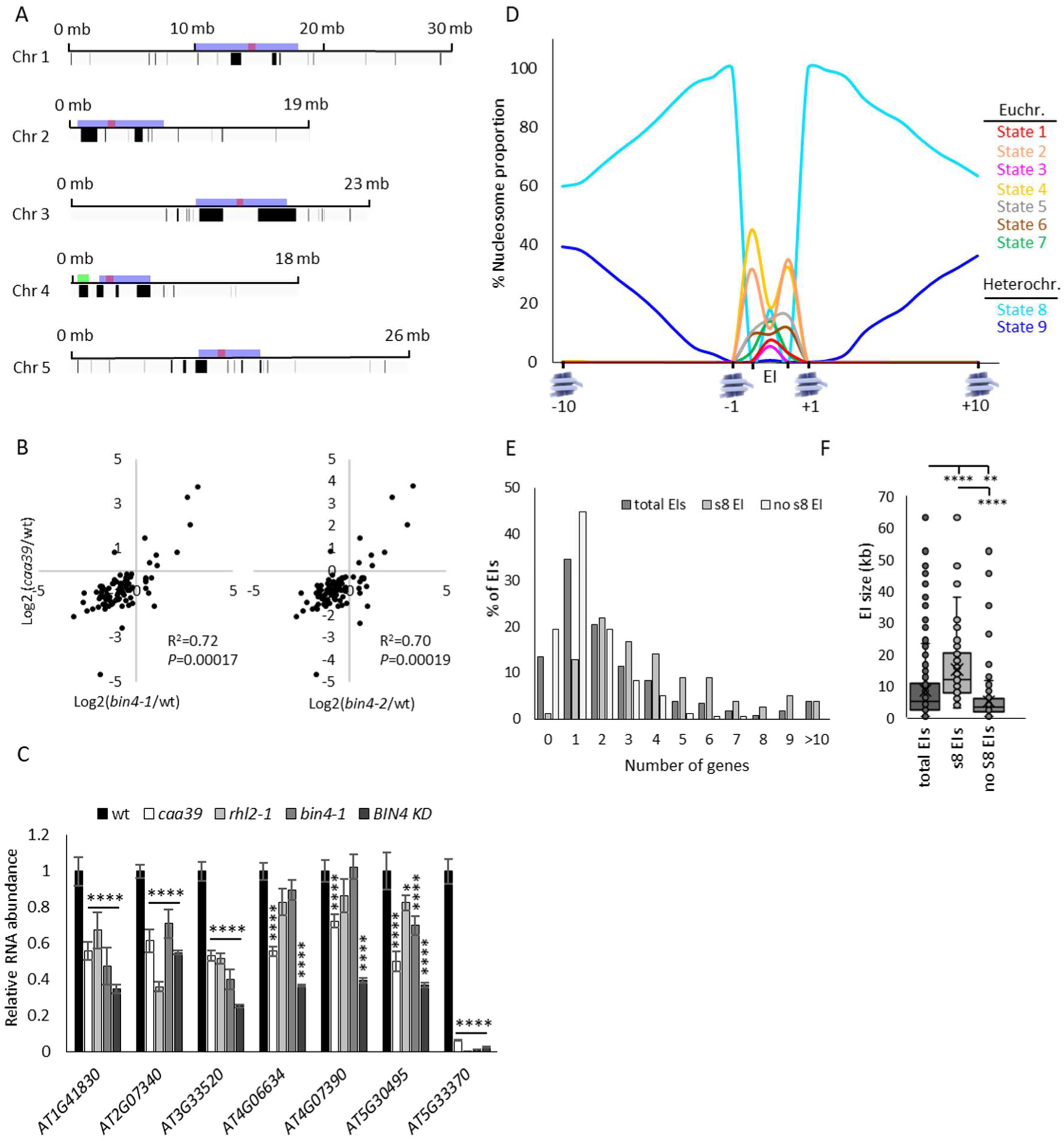
Genes in euchromatic islands within heterochromatic pericentromeric and chromosome 4 knob regions are repressed in Topo VI mutants. (A) Positional Gene Enrichment analysis of the top 500 most downregulated genes in *caa39*. The bed files corresponding to coordinates of the widest enriched regions (FDR < 0.05) were visualized with a genome browser. Black lines correspond to enriched regions, blue boxes correspond to pericentromeric regions as defined by Yelina et al. (2012), green box to the knob and red to centromeres. (B) Scatter plots and pearson correlations of differentially expressed (*P* < 0.05) pericentromeric genes in *caa39* and *bin4-1* or *bin4-2*. (C) RT-qPCR of selected pericentromeric genes in *caa39, rhl2-1*, *bin4-1* and *BIN4* KD at 6 days post-germination. Error bars: ± SEM of three biological replicates. **: *P* < 0.005; ***: *P* < 0.0005; ****: *P* < 0.0001 (two-way ANOVA, Dunnett’s test). (D) The proportion of each chromatin state was computed for ten nucleosomes on each side of the islands, along with the first nucleosome within each island. The proportions of euchromatic states within the islands correspond to the averaged proportions of all island nucleotides. EI: Euchromatic islands. (E) EI gene content in total, state 8-or no state 8-containing EIs. (F) EIs size distribution for total, state 8-or no state 8-containing EIs. **: *P* < 0.005; ****: *P* < 0.0001 (ANOVA, Tukey’s test).

### Downregulated genes within heterochromatic regions are localized in small euchromatic islands

We then asked whether the inverse expression patterns of genes and TEs in pericentromeric and *hk4S* regions in Topo VI mutants could be ascribed to their different chromatin landscapes. We first inspected the individual chromatin landscape of the seven downregulated pericentromeric genes confirmed by RT-qPCR (Fig 3C), using the nine chromatin states defined by Sequeira-Mendes *et al*. (Sequeira-Mendes *et al*, 2014). Euchromatin states 2, 1 and 3 characterize the proximal promoter, the transcriptional start site, and the start of coding sequence, respectively. The intragenic states 6 and 7 are characteristic of the transcriptional termination site and gene body of long transcribed genes, respectively. States 4 and 5 are highly enriched in H3K27me3 (a *Polycomb*-Repressive Complex 2 (PRC2)-based repressive histone modification) and are usually found in intergenic regions and PRC2-targeted genes. Lastly, the two types of heterochromatin states, 8 and 9, are enriched in H3K9me2, but in contrast with state 8, state 9 preferentially defines pericentromeric heterochromatin and is devoid of H3K27me3 (Sequeira-Mendes *et al*, 2014). Strikingly, all inspected loci share common features: a typical euchromatin context whose proximal environment is composed of heterochromatic state 8 and whose distal environment is of heterochromatic state 9 (Fig S6A). Overall, these observations suggested that *caa39* downregulated genes might be part of *bona fide* euchromatic islands (EIs).

To generate a comprehensive view of their structural features in the genome, we systematically investigated the pericentromeric and *hk4S* heterochromatic regions defined in Figure S6B. We designed a script to extract all EIs surrounded by chromatin states 8 and 9 and identified 232 EIs containing 540 EI genes this way, among which 6 correspond to unsequenced gaps (https://apps.araport.org/jbrowse) and were discarded in subsequent analyses (Appendix Table S3). Looking for EIs directly flanked by state 9 chromatin did not increase the number of EIs identified, showing that chromatin state 8 is always present in the proximal border (Fig 3D, Fig S6C). In contrast, the number of detected EIs started to decrease to 215 when considering at least 2 consecutive state 8 nucleosomes, indicating that 11 EIs have only one proximal nucleosome in state 8 (Fig S6C). With respect to state 9, the number of extracted islands began to drop from 4 consecutive nucleosomes (Fig S6C). To gain a better insight into the chromatin landscape of EIs, we analyzed the proportion of chromatin states covering EIs and ten nucleosomes on each side of EIs. This analysis revealed that EIs are mainly composed of chromatin states 1 to 7 with state 8 in 78 EIs (Fig 3D, Appendix Table S3). A majority of EIs are short (S-EIs) and contain 1 or 2 genes (Fig. 3E), and the 78 state 8-containing EIs were in average significantly longer (Fig 3F; Appendix Table S4) and contained more genes (Fig 3E, Appendix Table S3) than state 8-free EIs.

### Topo VI prevents the spreading of H3K9me2 into euchromatic islands

Given the general repression of EI genes and the local decrease of H3K9me2 at some TE loci without affecting the global level of H3K9me2 (Fig 1B), we hypothesized that EI gene repression might result from ectopic spreading of this silencing mark over EIs. This was first tested on several EI genes by ChIP-qPCR analysis of H3K9me2 levels in wt and *caa39*. H3K9me2 levels were very low, barely above background, in wt, consistent with the fairly high level of expression of these genes (Fig 4A). In contrast, a clear increase of H3K9me2 was observed in *caa39* (Fig 4A). Therefore, we further tested the spreading of H3K9me2 over EIs on a genome-wide scale by ChIP-seq analysis of H3K9me2 levels in wt and *caa39*, normalized to H3 levels in each line. Analysis of the wt profile of S-EIs (S-EIs<6kb) showed a chromatin landscape where H3K9me2 was barely detected, flanked by regions with high H3K9me2 levels (Fig 4B). Consistent with the minor decrease observed only on some TEs presented in Figure 2, the H3K9me2 level is globally not lower in EI-flanking sequences in *caa39* as compared to wt. In contrast, a clear increase was observed within S-EIs, which was highly significant all along S-EIs (Mann-Whitney test, *P* < 0.01), suggesting that Topo VI prevents H3K9me2 spreading across natural boundaries (Fig 4B). H3K9me2 spreading into large EIs (L-EIs) was also highly significant and particularly pronounced on L-EIs boundaries (Fig. 4C). Inspection of meta-profiles for each chromosome revealed highly significant H3K9me2 spreading into EIs (Fig S7A), that was observed for all replicates (Fig S7B). We then compared EIs to randomly generated euchromatin regions of the same length and observed that the most significant increase of H3K9me2 is located in EIs, although other chromosome arm regions may also gain slight amounts of H3K9me2 (Fig S8A and B). This shows that Topo VI exerts its H3K9me2 spreading inhibitory function mainly within heterochromatin regions.

**Figure 4.**
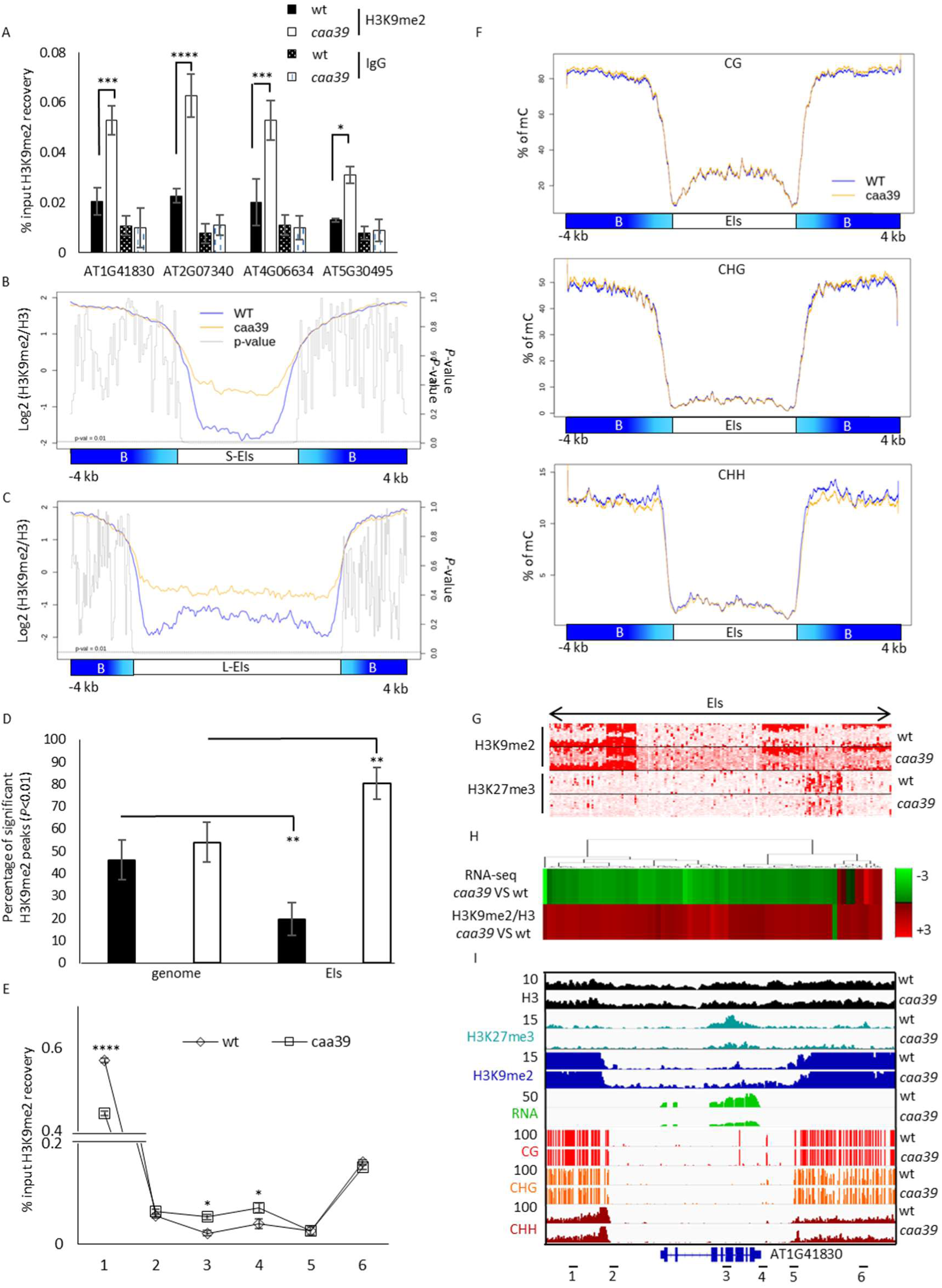
Topo VI prevents the invasion of euchromatic islands by H3K9me2. (A) ChIP-qPCR analysis of H3K9me2 at EI genes. Error bars: ± SEM of two biological replicates. *: *P* < 0.05; ***: *P* < 0.0005; ****: *P* < 0.0001 (two-way ANOVA, Fisher’s test). (B) Average distribution of H3-normalized H3K9me2 along short euchromatic gene islands and 4 kb flanking regions. Two (H3) and three (H3K9me2) independent biological replicates for each genotype were performed. *P*-value was computed for each aggregated point by using a Mann-Whitney test. (C) Same as (B) long EIs. (D) Bar chart showing the average percentage of down or up H3K9me2 peaks in the genome or in EIs. Error bars: ±SEM of three biological replicates. *: *P* < 0.05; **: *P* < 0.005; ***: *P* < 0.0005 (two-way ANOVA, Fisher’s test). (E) ChIP-qPCR validation of a single-gene island for H3K9me2. Error bars: ±SEM of three biological replicates. *: *P* < 0.05; ****: *P* < 0.0001 (two-way ANOVA, Fisher’s test). (F) Average distribution of methylated cytosine in CG, CHG and CHH contexts over EIs. Two independent replicates for each genotype were performed. (G) K-means linear clustering of H3K9me2 and H3K27me3 tag densities across EIs and their 900 nt flanking borders as revealed by seqMINER in *caa39* and wt. (H) Heatmap correlation clustering of EI genes with significant changes in RNA-seq and H3K9me2/H3 ChIP-seq. (I) Integrative Genomics Viewer screenshot of H3, H3K9me2, H3K27me3, RNA, CG, CHG and CHH methylation profiles in wt and *caa39* on a single gene-containing island. Each track is normalized against corresponding input samples (ChIP-seq) and by the sequencing depth. Numbers indicate the position of primers used in (E).

To further strengthen our analysis, we applied diffReps on each replicate individually (Shen *et al*, 2013) to confirm differential enrichment of H3K9me2 in EIs of *caa39* and wt (Appendix Table S5). This very stringent and not very sensitive analysis (essentially due to the fact that H3K9me2 peaks were barely detectable in wt and that H3K9me2 spreading does not appear to be sequence-specific) could still reveal increased levels of H3K9me2 in some EIs of *caa39*, but not all (90.4% in replicate 2, 70.5% in replicate 3, 21.7% in replicate 1; Appendix Table S3). The limited number of EIs identified this way in replicate 1 can be attributed to the fact that it was less deeply sequenced than replicates 2 and 3 (GSE103924). Despite that, there was a large overlap between replicates (Fig S7C). We then extracted the number of significantly (g test, *P* < 0.01) up or down H3K9me2 peaks in *caa39* versus wt, in each individual replicate (Appendix Table S5, sheet 6). A similar percentage of significantly up or down H3K9me2 peaks was observed at the genome scale by this method (Fig 4D), in agreement with the fact that the global level of H3K9me2 is not overtly affected in *caa39*. In contrast, the proportion of peaks was significantly shifted toward gains of H3K9me2 within EIs (Fig 4D), here again confirming the specific role of Topo VI in safeguarding EI genes from ectopic H3K9me2 marking.

Globally, these data show that Topo VI is required to prevent elevated H3K9me2 levels within EIs, presumably by preserving sharp boundaries between these insulated elements to avoid pervasive spreading of heterochromatin from flanking regions. We further confirmed such a barrier-like function on a S-EI containing a single gene, *At1g41830*, by ChIP-qPCR analysis of H3K9me2. Scanning of six different loci along this region (Fig 4E) in independent experiments confirmed the increased H3K9me2 levels within the island body, but also a decrease in one neighboring heterochromatic border (Fig 4E), similarly to what was observed in the ChIP-seq experiment replicates for this EI (Fig 4I and Fig S8C) and other inspected EIs (Fig S8C).

### EI gene repression is linked to H3K9me2 spreading

We then tried to evaluate the relative contribution of H3K9me2 increase to the down-regulation of EI genes, compared to other heterochromatic and repressive marks. Firstly, because H3K9me2 and non-CG (CHG and CHH) DNA methylation are strongly inter-dependent (Stroud *et al*, 2014), we tested whether H3K9me2 spreading into EIs might in turn, or reciprocally, affect DNA methylation. Whole-genome bisulfite sequencing revealed that DNA methylation levels in all sequence contexts were unaltered in *caa39* relative to wt in EIs (Fig 4F and I, Fig S8C). Differentially methylated region (DMR) analyses confirmed that there was no global significant increase of DNA methylation in EIs (Dataset S1). Surprisingly, despite CHGs being known to be methylated through a feedback loop with H3K9me2, DMR analyses rather revealed a decrease of CHG, with 51 hypo-CHG DMRs and 16 hyper-CHG DMRs observed in EIs of *caa39* (Dataset S1). Therefore, DNA methylation does not seem to contribute to the down-regulation of gene expression in the EIs and.

Secondly, because EI boundaries are enriched in H3K27me3-marked heterochromatin state 8, and that H3K27me3 is also found on the proximal promoter (chromatin state 2) and transcribed region of many euchromatic genes (chromatin state 5, silenced genes) (Sequeira-Mendes *et al*, 2014), we also performed a genome-wide H3K27me3 analysis by ChIP-seq. Interestingly, we observed a globally inverted tendency as compared to H3K9me2 profiles. Indeed, average H3K27me3 levels were locally and significantly decreased within EIs (Fig S9A-B). We further documented such a local decrease of H3K27me3 on the small EI containing *At1g41830*, by ChIP-qPCR (Fig S9C). Consistent with this, K-means linear clustering revealed that H3K27me3 decrease could not be generalized to all EIs, as it marks only a small subset of EIs in wt (Fig 4G). However, more global and significant H3K27me3 increased levels were observed in neighboring *caa39* heterochromatin (Fig S9A-B). Thus, on one hand, H3K27me3 does not seem to contribute to the global down-regulation of EI genes, and its local decrease on some EI genes might even counterbalance the effect of H3K9me2 increase in a few EIs. On the other hand, we observed more global and significant increases of H3K27me3 in heterochromatin regions at the border of EIs, suggesting that Topo VI, in addition to its role in preventing ectopic H3K9me2 marking within EIs, also prevents H3K27me3 to spread into heterochromatin borders of EIs.

Finally, we directly compared EI gene expression with H3K9me2 changes. A vast majority (90%) of EI genes that show significant changes (*P* < 0.05) in either RNA-seq or ChIP-seq analyses are repressed and possess enhanced levels of H3K9me2 (Fig 4H and I, Fig S10). Interestingly, the most repressed genes are short and tend to also possess the clearest increases of H3K9me2 (Fig 4H, Fig S10). Taken together, these results suggest that H3K9me2 increase plays a major role in the reduced expression of EI genes, although higher order chromatin structure, changes in other histone modification, and other indirect effects of the *caa39* mutation may contribute to gene expression changes.

### The Topo VI subunit BIN4 physically associates with MAT3

To gain knowledge on the molecular mechanism by which Topo VI contributes to the delimitation of chromatin boundaries, we used the BIN4 subunit as a bait to screen a yeast two hybrid (Y2H) cDNA library (Hybrigenics). A strong interaction with the Topo VI subunit RHL1 was detected, thereby demonstrating the reliability of the screening procedure (Appendix Table S6). Among the eleven additional interacting partners, three proteins belong to the S-AdenosylMethionine (SAM) biosynthesis pathway, the universal methyl group donor (Zhang, 2005). The first one, 5-methylthioribose-1-phosphate isomerase (MTI1, AT2G05830), is involved in the methionine salvage pathway whereas the two others, Methionine Synthase 1 (MS1, AT5G17920) and Methionine Adenosyltransferase 3 (MAT3, AT2G36880) are the ultimate enzymes of the SAM cycle. In order to identify BIN4-interacting proteins *in planta*, we also performed a CoIP-MS experiment using the *Arabidopsis mid-1 35S:BIN4/MID-YFP* line, which consists of the *mid-1* allelic mutant of *BIN4* complemented with YFP-tagged BIN4/MID (Kirik *et al*, 2007), and a wt line as control. To exclude nonspecific proteins, we discarded proteins that were not detected in both BIN4/MID-YFP CoIP-MS replicates, as well as chloroplastic, mitochondrial and peroxysomal proteins. The presence of the RHL1 and TOP6B Topo VI subunits in BIN4/MID-YFP CoIPs validated our procedure (Appendix Table S7). Remarkably, MAT3 co-immunoprecipitated strongly with BIN4. We further investigated the genetic and biochemical interactions between BIN4 and enzymes of the SAM cycle, particularly the very last enzyme MAT3, using Bimolecular Fluorescence Complementation (BiFC). We confirmed the BIN4-MAT3 and BIN4-MS1 interactions in nuclei of transiently agro-transformed *Nicotiana benthamiana* mesophyll cells (Fig 5A).

**Figure 5.**
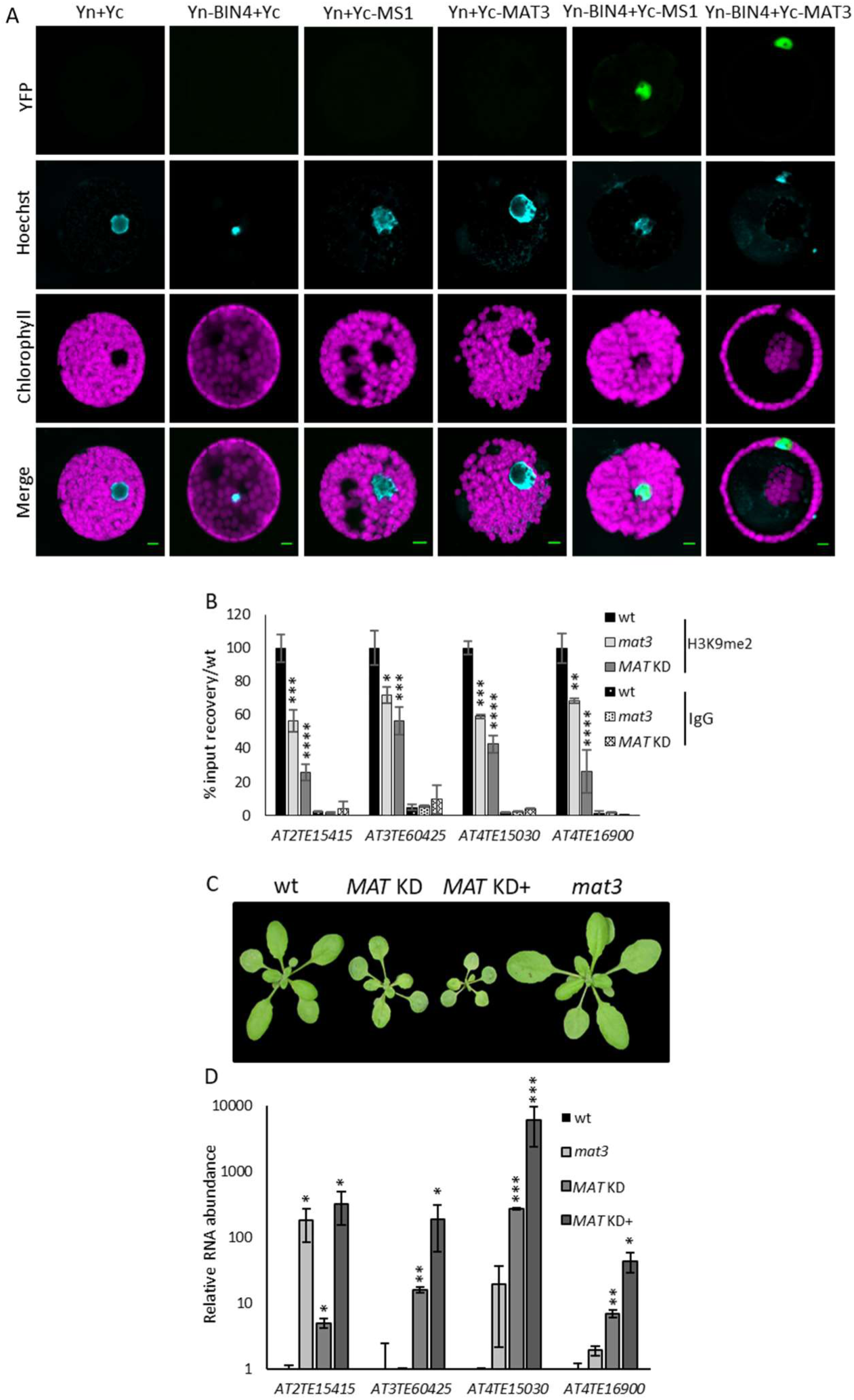
MAT3 directly interacts with Topo VI and is required for H3K9me2 deposition on heterochromatic loci. (A) Protoplasts from transiently agrotransformed *N. benthamiana* leaves expressing different combination of BiFC vectors, as indicated. Nuclei were stained with Hoechst 33342. (B) Chromatin of 3 week-old wt or *MAT* KD (here a mixed pool of *MATs* silenced plants with strong or weak phenotypes) rosette leaves, or 6 day-old wt or *mat3* cotyledon nuclei was immunoprecipitated with anti-H3K9me2 antibodies and the recovery of TEs known to be reactivated in *caa39* was assessed by qPCR. The result is shown as a percentage of recovery normalized against wt. Error bars: ± SEM of two biological replicates. *: *P* < 0.05; **: *P* < 0.005; ***: *P* < 0.0005; ****: *P* < 0.0001 (two-way ANOVA, Dunnett’s test). (C) Representative photographs of 4 week-old wt, *MATs* silenced plants with stronger (*MAT* KD+) and weaker (*MAT* KD) developmental phenotypes, and the *mat3* mutant. (D) RT-qPCR analysis of TE transcript abundance in indicated genotypes. Error bars: ± SEM of three biological replicates. *: *P* < 0.05; **: *P* < 0.005; ***: *P* < 0.0005 (Student’s *t*-test).

### MAT3 is required for H3K9me2 deposition

Given that MAT enzymes synthesize the SAM required for DNA and histone methylation, and that Topo VI is required for proper distribution of H3K9me2 throughout EI-containing heterochromatic regions, we hypothesized that disruption of MAT3 affects H3K9me2 deposition. To address this question, we used a recently characterized knock-down line in which *MAT3* 3’-UTR is interrupted by a T-DNA (Chen *et al*, 2016), generating strongly reduced but still detectable transcripts levels (Fig S11A). We measured H3K9me2 levels at the four TEs strongly de-repressed in Topo VI mutant plants (Fig 2) by ChIP-qPCR and found decreased levels of H3K9me2 (Fig 5B). This modest decrease of H3K9me2 might be explained by the hypomorphic nature of the mutation or by a functional redundancy between MAT isoforms that share over 85% amino acid sequence identity (Fig S11B). To test this hypothesis, we took advantage of a homozygous, mono-insertional, co-suppressor transgenic line obtained during the process of generating *MAT3-YFP* overexpressors, that we referred to as *MAT* KD (Fig 5C, Fig S11C). Owing to their high DNA sequence identity (Fig S11D), all *MAT* genes are downregulated in this line (Fig S11E). In addition, the stochastic silencing of *MATs* gives rise to different phenotype severities: mildly affected *MAT* KD plants (Fig 5C) that accumulate less *MAT*s transcripts than wt (Fig S11E) and present a more severe phenotype than *mat3* hypomorphic mutant plants (Fig 5C); and strongly affected *MAT* KD+ sister plants (Fig 5C) that accumulate even less *MAT* transcripts (Fig S11E). As anticipated, the H3K9me2 decrease was even more pronounced in *MAT* KD plants than in *mat3* (Fig 5B). These results suggest that MAT isoforms possibly have additive roles in H3K9 dimethylation. Given the decrease of H3K9me2 in MAT-deficient plants, we then determined the extent of TE reactivation in *mat3* and *MAT* KD by RT-qPCR analysis of the same four TEs. We observed increased levels of TE transcripts in *mat3* (Fig 5D). This increase was generally more pronounced in *MAT* KD plants, particularly in *MAT* KD+ plants (Fig 5D). In contrast, we did not observe any significant effect on EI gene expression (Fig S11F).

### Topo VI favors MAT enrichment at some heterochromatin borders and depletion from euchromatic islands

Collectively, our results suggest that Topo VI and MAT3 could act together in maintaining sharp chromatin boundaries by influencing H3K9me2 deposition. We therefore used a newly developed anti-MAT antibody (Agrisera, Vännäs, Sweden) to test for MAT enrichment at specific loci and a putative Topo VI dependency. First, the specificity of this antibody was validated by immunoblot analysis of protein extracts from wt, *MAT* KD and *MAT3-YFP* overexpressing lines (Fig S12A). We then performed anti-MATs ChIPs on wt and *caa39* plants to measure the recovery of TEs that are reactivated upon Topo VI or MATs loss of functions. Interestingly, MATs were enriched on all the tested TEs in wt and less in *caa39*, as compared to an IgG negative control (Fig 6A). To specifically evaluate the implication of the BIN4/MID-associated MAT3 isoform, we undertook an anti-GFP ChIP analysis of two independent MAT3-YFP expressing lines. TEs reactivated in *MAT* KD and in Topo VI mutant plants were also specifically enriched in the GFP-pulled down chromatin (Fig S12B). These results indicate that Topo VI is required for the association of MAT3 with heterochromatic elements to enable local deposition of H3K9me2.

**Figure 6.**
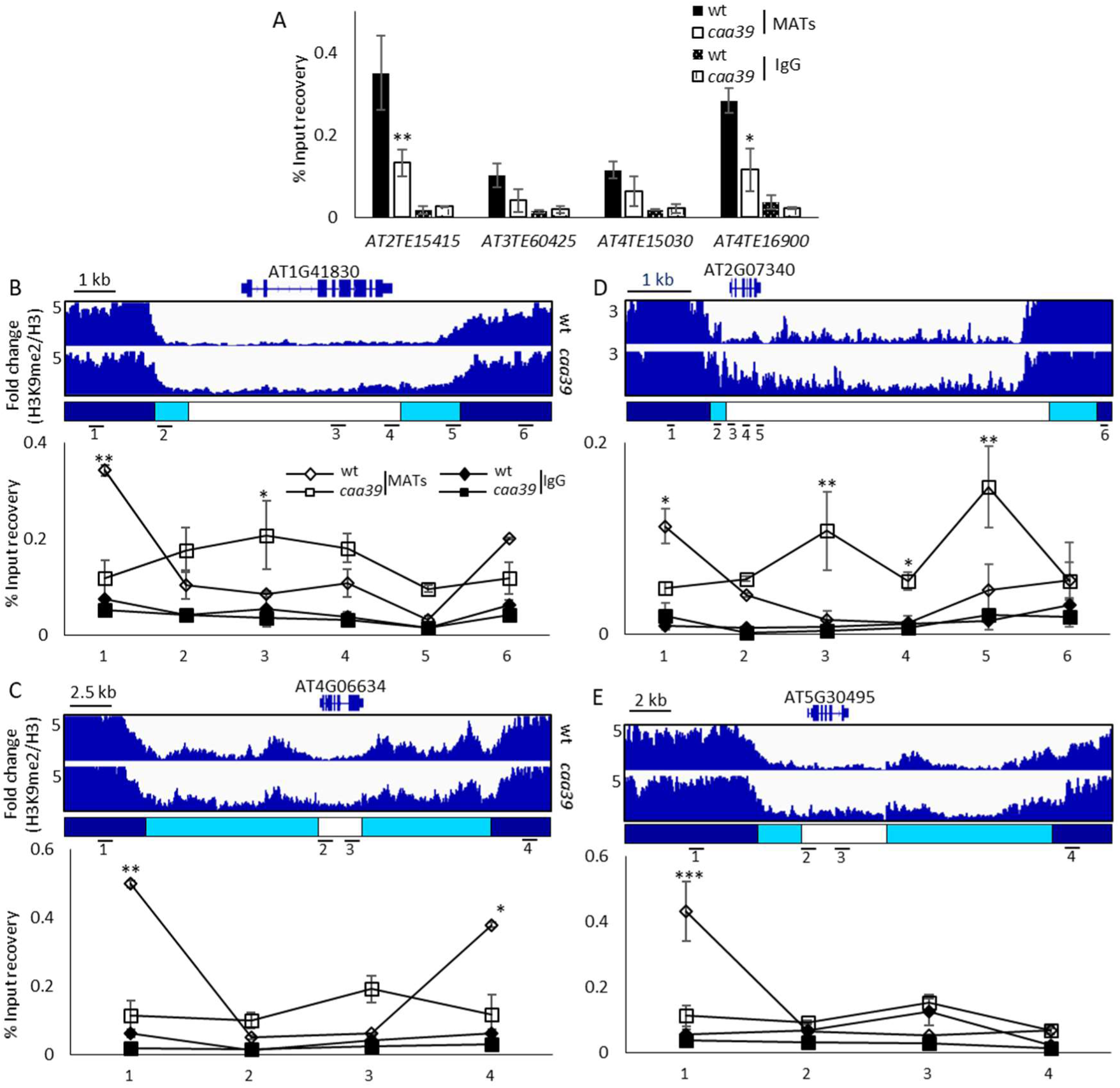
Topo VI is required for MAT enrichment at heterochromatin borders and exclusion from euchromatic islands. (A) Chromatin of 6 day-old wt or *caa39* cotyledon nuclei was immunoprecipitated with anti-MATs antibodies and the recovery of TEs reactivated in *MATs* silenced plants and *caa39* was measured by qPCR. Error bars: ± SEM of three biological replicates. (B-E) Top panels: Integrative Genomics Viewer screenshots of H3-and sequencing depth-normalized H3K9me2 profiles, and locations of primers used in bottom panels. Bottom panels: Same as (A) on EIs containing repressed genes in *caa39* such as *At1g41830* (B), *At4g06634* (C), *At2g07340* (D) and *At5g30495* (E). Error bars: ± SEM of two biological replicates. *: *P* < 0.05; **: *P* < 0.005; ***: *P* < 0.0005 (Student’s *t*-test).

To test whether Topo VI might also influence the enrichment of MATs at EIs, we conducted an anti-MATs ChIP experiment on *caa39* and wt plants and probed the same EI as in Figures 4E and I, which shows increased H3K9me2 levels within the island body but decreased levels in one heterochromatic border in *caa39* (Fig 4E and I, Fig 6B, top panel). We detected a significant decrease of MAT occupancy in this border in *caa39* (Fig 6B, bottom panel, probe 1) and, conversely, enhanced MAT occupancy in the island body in *caa39* compared to wt (Fig 6B, bottom panel, probes 3-4). We analyzed three other S-EIs that show decreased gene expression levels (Fig 3C), increased internal H3K9me2 levels (Fig 4C and S8C) and contain a single gene (*At4g06634*’s and *At5g30495*’s EIs) or two genes (*At2g07340*’s EIs). We generally observed a decrease of MAT occupancy at one or both borders and enhanced MAT occupancy in the island bodies of *caa39* (Fig 6C-E). These results show that the loss of Topo VI leads to MAT redistribution over EIs, which positively correlates with H3K9me2 redistribution and EIs heterochromatinization in *caa39*.

## Discussion

The *Arabidopsis* epigenome is largely indexed by discrete chromatin signatures usually corresponding to single genetic element (e.g., a gene or a TE) (Roudier *et al*, 2011; Sequeira-Mendes *et al*, 2014). However, despite this, in *Arabidopsis* DNA methylation has a known tendency to spread away from many TEs (Ahmed *et al*, 2011), and a few other studies have reported the existence of heterochromatin spreading in plants (Eichten *et al*, 2012; Lang *et al*, 2015; Saze *et al*, 2008). Yet, the mechanisms that repress heterochromatin spreading, hence safeguarding EIs, are poorly understood in plants. Our study confirmed the existence of an insulator-like mechanism that preserves EIs and unveiled the role played by the Topo VI complex in this process. We first provide evidence that Topo VI is required to preserve the euchromatic nature and transcriptional activity of gene islands within pericentromeric and chromosome 4 knob heterochromatic regions. Indeed, the most remarkable effect of the *caa39* mutation was the specific misregulation of pericentromeric elements, with a general downregulation of EI genes and, inversely, a reactivation of heterochromatic TEs. We confirmed this peculiar expression pattern in several amorphic and hypomorphic mutants of the Topo VI complex. Surprisingly, EI gene downregulation is more pronounced in the hypomorphic mutants *caa39* and *BIN4* KD than in the corresponding null Topo VI mutants that display more severe growth defects. Taking advantage of uncoupled growth defects and gene expression changes in the hypomorphic *caa39* allele, we were able to show that the repression of EI genes is correlated with the invasion of EIs by H3K9me2, indicating that Topo VI is required for preventing H3K9me2 spreading, here referred to as a boundary function. The level of H3K9me2 in EIs of *caa39* remain, however, significantly lower than in canonical heterochromatin. This might result from histone H3 demethylation, a process that is likely still active in *caa39* as we did not observe any alteration of expression of key demethylation genes such as *IBM1*. Alternatively, gain of H3K9me2 could be the outcome of a subset of cells in which *caa39* mutation has a strong effect on H3K9me2 boundaries.

Unexpectedly, WGBS identified no global increase of cytosine methylation over EIs, but rather a decrease of CHG methylation, despite CHGs being known to be methylated through a feedback loop with H3K9me2. Recent studies report that, under specific circumstances, increased H3K9me2 levels do not necessarily induce increased CHG or CHH methylation, and *vice versa*. For instance, the AT-hook protein AHL10 ectopically recruits H3K9me2 to small, AT-rich TEs without coincidental increases in DNA methylation (Jiang *et al*, 2017). Another study reports that expression of *AtCMT3* in *Eutrema Salsugineum*, a Brassicaceae that has lost *CMT3* and gene body methylation, indeed induce *de novo* gene body methylation in CHG, CHH and CG contexts, but does not induce gains in H3K9me2. Interestingly, hyper-CHG methylation in gene bodies was not correlated with consistent changes in gene expression (Wendte *et al*, 2019). More generally, the interplay between H3K9me2 and DNA methylation might be regulated by the kinetics of activity of both pathways. Indeed, heterochromatin is known to inhibit and exclude RdDM (Zemach *et al*, 2013; Schoft *et al*, 2009; Gent *et al*, 2014). Thus, it is plausible that H3K9me2 spreading over EI genes is sufficient to induce transcriptional silencing without the involvement of DNA methylation in a context of lack of Topo VI. Nonetheless, we have no indication to rule out a possible role of DNA methylation and/or ncRNAs in the delineation of sharp chromatin boundaries as EIs. Interestingly, RNA Pol V-produced ncRNAs also play a role in the determination of heterochromatic identity in *Arabidopsis* but are very unlikely involved in the inhibition of heterochromatin spreading (Böhmdorfer *et al*, 2016). Inversely, RNA-directed DNA methylation (RdDM) appears to be required to inhibit the spreading of euchromatin into heterochromatin in maize (Li *et al*, 2015). Nevertheless, TEs can provide insulator sequences in animals (Wang *et al*, 2015b) and the activity of CTCF is known to be regulated by ncRNAs (Cohen & Jia, 2015). In fission yeast, the non-coding RNA (ncRNA) *BORDERLINE* prevents the spreading of heterochromatin to euchromatin by evicting the H3K9me reader and chromatin barrier protein Swi6/HP1 from euchromatin (Keller *et al*, 2013).

The disorganization of chromocenters and the loss of pericentromeric TE silencing observed in Topo VI mutants might result from the loss of this boundary function, and likely also by the loss of an distinct architectural function of Topo VI in heterochromatin condensation. Indeed, given that no obvious loss of DNA methylation and only a partial decrease of H3K9me2 over TEs were observed in Topo VI mutant plants, the reactivation of TEs is unlikely to be solely explained by decreased levels of these marks, but rather by a combined loosening of their heterochromatic nature and of their higher order organization. It is worth noting that in animals, insulator proteins can participate in several distinct processes. For instance CTCF acts locally as a chromatin barrier and more globally on the formation of topologically associating domains (Lu *et al*, 2016). Furthermore, the co-occurrence of EI invasion by H3K9me2 and the increase of H3K27me3 within heterochromatin supports a dual breaking model of the *Arabidopsis* Topo VI boundary function. Alternatively, heterochromatin spreading might itself perturb H3K27me3 deposition by PRC2 and/or erasing by trithorax group (trxG) proteins. This second hypothesis is supported by similar observations made in other organisms, where the loss of Swi6/HP1 leads to H3K9me2 spreading across natural constitutive heterochromatin boundaries in fission yeast (Stunnenberg *et al*, 2015), and alters H3K27me3 deposition on facultative heterochromatin in *Neurospora crassa* (Jamieson *et al*, 2016).

Notably, other plant GHKL ATPases such as the MORC proteins are also required for TE silencing and chromocenter formation without strongly impacting H3K9me2 and DNA methylation levels (Moissiard *et al*, 2012; Lorkovi *et al*, 2012; Brabbs *et al*, 2013). Although *atmorc1* and *atmorc6* mutants appear to be less dramatically affected than c*aa39*, Topo VI might act downstream of DNA methylation in a similar manner to what Moissiard *et al*. reported for MORC proteins (Moissiard *et al*, 2012). Interestingly, human CTCF has been shown to interact with Topo IIβ (Witcher & Emerson, 2009; Yusufzai *et al*, 2004) and appears to be part of a protein interaction network that also contains MORC2 and members of the cohesin complex in Hela cells (Uusküla-Reimand *et al*, 2016). Intriguingly, plant and human recombinant MORC proteins seem to display a type II topoisomerase-like activity, which requires additional factors for full activity. However, the possibility that topoisomerases co-purify with MORC proteins cannot be completely ruled out (Manohar *et al*, 2017). Collectively, our study on plant Topo VI extends the role of topoisomerase function to chromatin, which is an emerging theme in a wide range of organisms.

Although we cannot rule out the implication of other MATs in the definition of boundaries and/or EI identity, the finding that the Topo VI BIN4 subunit directly interacts with MAT3 and is required for MAT3 enrichment at TE loci and depletion from euchromatic islands, argues for a local role of MAT3 on chromatin, in addition to its general role in SAM synthesis. Interestingly, a homologue of MAT3 in mammals, MATIIα, also directly supplies SAM in the close vicinity of oncogenes to allow transcriptional repression and H3K9me2 deposition (Katoh *et al*, 2011; Kera *et al*, 2013). The mouse MATIIα has further been found to interact with Topoisomerase IIα, a type IIA topoisomerase whose C-terminal regulatory domain possesses sequence similarity with the BIN4 subunit of the plant Topo VI complex (Breuer *et al*, 2007). Hence, although this requires testing in other organisms, an interaction between MAT enzymes and type II topoisomerases might be evolutionary conserved. Targeting of MATs to specific chromatin regions by topoisomerases might be a way to couple SAM synthesis and availability *in situ*, possibly for DNA or histone methylation. An analogous system might also exist between MAT, histone methyltransferases and type I topoisomerases in plants. Indeed, Xuemei Chen’s group has recently reported that Topo Iα, in addition to being involved in TE silencing through RdDM and H3K9 dimethylation (Dinh *et al*, 2014), is also required for *Polycomb*-mediated gene regulation (Liu *et al*, 2014). Interestingly, by mining mass spectrometry data of immuno-purified CLF and ALP1 *PcG* subunits (Liang *et al*, 2015), residual levels of MAT3 and more abundant levels of MAT4 could be identified (data not shown). In plants, the existence of an insulator-like function that would partition chromatin into different functional domains has long been questioned (Wang *et al*, 2015a; Liu *et al*, 2017; Vergara & Gutierrez, 2017). We show that Topo VI participates to such a function by preventing the spreading of the heterochromatic mark H3K9me2 into neighboring euchromatin islands. Our results suggest that the prevention of heterochromatin spreading relies upon Topo VI-dependent targeting of MAT3 to heterochromatin, and exclusion from euchromatic islands. Future studies might allow the identification of direct molecular links between MAT proteins and methyltransferases involved in DNA, H3K9 and/or H3K27 methylation for fine-tuning the establishment of sharp transitions in chromatin identity along the genome.

## Materials and Methods

### Cloning and plasmids

cDNA containing the full-length *BIN4.3* (*AT5G24630.3*), *MAT3* (*AT2G36880*) and *MS1* (*AT5G17920*) open reading frames were amplified by RT-PCR from *Arabidopsis thaliana* ecotype Col-0 and with primers listed in Additional file 8. The PCR products were introduced by BP recombination into the pDONR207 entry vector (Invitrogen) and sequenced to ensure the absence of mutations. These entry clones, with or without a stop codon, were then introduced into destination vectors using LR recombination and sequenced again to confirm that the fusions were in frame. To generate transgenic lines, the entry clones were transferred to pEarleyGate 102 (BIN4-CFP-HA) or pEarleyGate 101 (MAT3-YFP-HA) binary vectors (Earley *et al*, 2006). For BiFC assays, the entry clones were transferred to the four pBiFP vectors (Azimzadeh *et al*, 2008). All binary vectors for transient or stable plant transformation were then transferred to *Agrobacterium tumefaciens* strain C58C1 RifR (pMP90) by electroporation.

### Plant material and growth conditions

All lines are in the Col-0 ecotype. *ddm1-8* (SALK_000590C) was ordered from the NASC collection. The *rhl2-1*, *bin4-1, caa39* and *mat3* mutants are already described (Breuer *et al*, 2007; Chen *et al*, 2016; Simkova *et al*, 2012; Sugimoto-Shirasu *et al*, 2002). The *MAT3-YFP* and *BIN4-CFP-HA* transgenic lines were generated using the floral dip method (Zhang *et al*, 2006) and single insertion homozygous lines were selected. In all experiments except those using the *MAT* KD line, plants were cultivated on half strength MS/Agar medium with 16 h light/8 h dark cycles for six days at 80-90 µmol photons m^−2^ s^−1^ light intensity. For other experiments, plants were cultivated in soil for three to four weeks under the same light regime as stated above.

### DNA preparation, Chop-qPCR and Anti-5-meC ELISA assay

Genomic DNA was prepared using the NucleoSpin® Plant II Midi kit (Macherey-Nagel). For each reaction (either with McrBC, HpaII, MspI or HaeIII), 200 ng DNA was used in a 20 µL final reaction volume. 5 units of MspI, HpaII or HaeIII per 100 ng DNA were used during 1 h at 37 °C. For McrBC digestion, 1 unit per 100 ng DNA was used for an overnight digestion at 37 °C. One nanogram of undigested or digested samples was used for qPCR. Undigested samples were treated at the same temperature and for the same time but without enzyme. The result is expressed as 2^-(Ct^_digested_-^Ct^_undigested_^)^. Global methylation of cytosines was assessed using the MethylFlash global DNA methylation ELISA easy kit following the manufacturer’s instructions (Epigentek).

### Yeast Two-hybrid Screen

The yeast two-hybrid screen was performed by Hybrigenics using the *Arabidopsis* RP1 library. The full-length *BIN4* cDNA (*AT5G24630.3/4*) was used as bait.

### Co-immunoprecipitation-MS

500 mg of ten-day-old wt and *mid1 MID-YFP* seedlings were harvested and ground in liquid nitrogen, mixed with 2 mL of RIPA buffer (10 mM TrisHCl pH 7.5, 150 mM NaCl, 0.5 mM EDTA, 0.1% SDS, 1% Triton X-100, 1% sodium deoxycholate, protease inhibitors). Samples were incubated for 30 min in ice and gently resuspended every 10 min, centrifuged for 15 min, 16,100 x *g*, 4 °C. For bead preparation, rabbit monoclonal anti-BIN4 (Breuer *et al*, 2007) and rabbit polyclonal anti-GFP antibodies (Thermo-Fisher, Cat. No. A-11122) were added in equal quantity (16 µg each) to 80 µL of Dynabeads® protein A (Thermo-Fisher, Cat. No. 10001D). Antibodies were incubated with beads overnight at 4 °C before crosslinking with DMP following the manufacturer’s protocol (Abcam). Then, antibody-conjugated beads were incubated with protein extracts for 2 h at 4 °C, and washed three times in immunoprecipitation buffer (1 mM EDTA, 10% glycerol, 75 mM NaCl, 0.05% SDS, 100 mM TrisHCl pH 7.4, 0.1% Triton X-100). Proteins were eluted by incubating at 70 °C in Laemmli buffer and the eluates were briefly migrated on SDS-PAGE before MS analysis.

### Western blot

For nuclear proteins, nuclei were prepared as described in the ChIP protocol below. The nuclei pellet was resuspended in nuclei lysis buffer and proteins were quantified using the bicinchoninic acid assay. 60 μg of nuclear proteins was separated on 12% SDS-PAGE and transferred to PVDF membranes. For total protein extracts, tissue powder was resuspended in SDS loading buffer (v/v), centrifuged 16,100 x *g*, 10 min, 4 °C, and boiled 5 min. BIN4, H3K9me2 (Abcam ab1220, lot: GR244373-6), H3 (Abcam ab1791, lot: GR178101-1), MAT (Agrisera AS16 3148) and GFP (Roche 11814460001, lot: 12600500) antibodies were diluted 1:3000, 1:2000, 1:1000, 1:5000, 1:3000 and 1:3000 in TBST/milk 0.5%, respectively. After blocking 1 h in 5% milk, membranes were incubated with the primary antibody for 1 h, washed 5 times with TBST and incubated with secondary HRP-coupled anti-mouse or anti-rabbit antibodies at 1:20000 during 1 h. Membranes were washed 5 times and revealed with Immobilon western chemiluminescent HRP substrate (Millipore).

### RNA extraction, RT-qPCR and microarrays

RNA was extracted using Trizol following the manufacturer’s instructions. The integrity of RNA was verified by migrating 1 µg RNA on a 1.5% agarose gel. Then, 500 ng RNA was DNAse I-treated (ThermoFischer Scientific) and submitted to reverse transcription (Takara PrimeScript™ RT). Before qPCR, the absence of genomic DNA and equal RNA loading per sample was verified using semi-quantitative RT-PCR against *ACT2* (see Appendix Table S8 for a complete primer list). cDNAs were diluted twice and 1 µL was used per qPCR reaction in a final volume of 15 µL. *PP2A* (*AT1G13320*) and *PRF1 (AT2G19760*) were used as control genes in every RT-qPCR experiment. Affymetrix Arabidopsis ATH1 GeneChips were used to analyse *bin4-1* and *bin4-2* mutant lines. Raw data processing was performed using the RMAExpress software (http://rmaexpress.bmbolstad.com/). Data were normalized by using background adjusting and quantile normalization.

### RNA-seq library preparation and sequencing

Three independent biological replicates were produced. For each biological repetition, RNA samples were obtained by pooling RNAs from more than 100 plants. Aerials parts were collected on plants at 1.00 developmental growth stages (Boyes *et al*, 2001), cultivated as described above. Total RNA was extracted using RNeasy kit (Qiagen®, Hilden, Germany) according to the supplier’s instructions. RNA-seq experiment was carried out at plateform POPS, transcriptOmic Plateform of institute of Plant Sciences - Paris-Saclay, thanks to IG-CNS Illumina Hiseq2000 privileged access to perform paired-end 100bp sequencing, on RNA-seq libraries constructed with the TruSeq_Stranded_mRNA_SamplePrep_Guide_15031047_D protocol (Illumina®, California, U.S.A.). The RNA-seq samples have been sequenced in paired-end (PE) with a sizing of 260 bp and a read length of 100 bases. Six samples by lane of Hiseq2000 using individual bar-coded adapters and giving approximately 30 million of PE reads by sample were generated.

### RNA-seq bioinformatic treatment and analysis

To facilitate comparisons, each RNA-Seq sample followed the same workflow from trimming to transcript abundance quantification, as follows. Read preprocessing criteria included trimming of library adapters and performing quality control checks using FastQC (v0.10.1). The raw data (fastq) were trimmed for Phred Quality Score > 20, read length > 30 bases and sort by Trimming Modified homemade fastx_toolkit-0.0.13.2 software for rRNA sequences. Bowtie v2 (Langmead & Salzberg, 2012) was used to align reads against the *Arabidopsis thaliana* transcriptome (with --local option). The 33602 mRNAs were extracted from TAIRv10 database (Lamesch *et al*, 2012) with one isoform per gene to avoid multi-hits. The abundance of mRNAs was calculated by a local script which parses SAM files and counts only paired-end reads for which both reads map unambiguously to the same gene. According to these rules, around 98% of PE reads aligned to transcripts for each sample. Gene expression analysis was performed using likelihood ratio test for a negative binomial generalized linear model where the dispersion was estimated by the edgeR method (v1.12.0, (McCarthy *et al*, 2012)) in the statistical software ‘R’ (v2.15.0, (R Development Core Team, 2017)). TEtranscripts (Jin *et al*, 2015) and edgeR were used to discover differentially expressed transposable elements in *caa39* compared to wt. The reference genome was *Arabidopsis_thaliana* TAIR10.31.gtf and the TE reference was TAIR10_TE.gtf. TEtranscripts options: multi-mapper mode and SAM files.

### Whole-genome bisulfite sequencing and DNA methylation analysis

Seeds of wt and *caa39* were sowed with an interval of one week to generate biological replicates. For each replicate, DNA samples were obtained from 400 mg of fresh aerial parts of 6 day-old plants (developmental growth stages (Boyes *et al*, 2001)) crushed in liquid nitrogen. Genomic DNA was prepared using the NucleoSpin® Plant II Midi kit (Macherey-Nagel), with the addition of Poly(vinylpyrrolidone) to the powdered sample. Libraries were prepared and sequenced at the BGI (China). Quality control was performed on sequencing files using FASTQC (v0.11.5) and adapter trimming was performed at BGI. Methylation analysis has been performed with Bismark (v0.20.0) (Krueger & Andrews, 2011) using default parameters, except –CX option, to get data for all the three contexts (CG, CHH and CHG). The bedgraph output file has been filtered into 3 files, one for each context. DMRfinder (Gaspar & Hart, 2017) was used to identify differentially methylated regions and the “combine_CpG_sites.py” script was used to cluster CpG (and other context) sites into regions with data_bismark.cov files as input files and default parameters. findDMRs.r was used with default parameters to conduct pairwise tests of sample groups in order to identify DMRs.

### Chromatin Immunoprecipitation

Two hundred 6-day-old seedlings were used per biological replicates. Seedlings were quickly harvested using a cat hair comb, root material was cut-off and the remaining photosynthetic material was fixed with 1% formaldehyde for 10 min. Crosslinking was quenched by addition of glycine at 125 mM final concentration and 5 min incubation, washed twice in ddH2O and the excess water was drained using kimwipes. Plant material was then ground in liquid nitrogen, resuspended in extraction buffer 1 (0.4 M Sucrose, 10 mM Tris pH 8, 10 mM MgCl_2_, 5 mM β-mercaptoethanol and protease inhibitor), filtered with four layers of miracloth and centrifuged 20 min at 3,300 x *g*, 4 °C. The pellet was resuspended in extraction buffer 2 (0.25 M Sucrose, 1% Triton X100, 10 mM Tris pH 8, 10 mM MgCl_2_, 5 mM β-mercaptoethanol and protease inhibitor) and centrifuged 10 min, 16,100 x *g*, 4 °C. The pellet was resuspended in 300 µL extraction buffer 2 and layered on top of 300 µL extraction buffer 3 (1.75 M Sucrose, 0.15% Triton X100, 10 mM Tris pH 8, 2 mM MgCl_2_, 5 mM β-mercaptoethanol and protease inhibitor) and centrifuged at 16,100 x *g* for 1 h, 4 °C. The resulting pellet was resuspended in 130 µL of nuclear lysis buffer (50 mM Tris pH 8, 10 mM EDTA, 1% SDS and protease inhibitor) and sheared using a Bioruptor for 10 cycles, high settings, 30 sec ON/60 sec OFF, twice. Then, the sheared chromatin was diluted ten times in ChIP dilution buffer (1.1% Triton X100, 16.7 mM Tris pH 8, 1.2 mM EDTA, 167 mM NaCl and protease inhibitor). In parallel, 10 µL protein G Dynabeads® were washed twice in ChIP dilution buffer and incubated either with 2 µg of mouse IgG (Sigma), anti-H3K27me3 (Abcam ab6002, lot: GR275911-2), anti-H3K9me2, anti-MAT or anti-GFP during 2 h at 4 °C under gentle agitation. Antibody-conjugated beads were washed twice with 100 µL ChIP dilution buffer and 400 µL of input was submitted to IP for 4 h or at 4 °C under gentle agitation. Immunocomplexes were washed twice with each following buffer, 5 min, 4 °C, gentle agitation, in that order: low salt wash buffer (150 mM NaCl, 0.1% SDS, 1% Triton X100, 2 mM EDTA, 20 mM Tris pH 8), high salt wash buffer (500 mM NaCl, 0.1% SDS, 1% Triton X100, 2 mM EDTA, 20 mM Tris pH 8), LiCl wash buffer (250 mM LiCl, 1% Igepal CA-630, 1% sodium deoxycholate, 1 mM EDTA, 10 mM Tris pH 8), TE buffer (10 mM Tris pH 8, 1 mM EDTA). Immunocomplexes were then eluted twice with 100 µL elution buffer (100 mM NaHCO3, 1% SDS), 15 min at 65 °C and agitation. The 200 µL eluate was reversed crosslinked by adding 16 µL of 2.5 M NaCl and incubating at 65°C overnight and DNA was purified using phenol/chloroform extraction after RNase A and proteinase K treatment. A 40 µL aliquot of input was reversed crosslinked and purified in parallel. DNA was quantified using Qubit® dsDNA HS assay kit (ThermoFischer Scientific) and equal amounts of input or IPed DNA were used per qPCR reaction.

### ChIP-seq analysis

Each ChIP biological replicate was prepared by pooling two independent IP eluates. ChIP-seq librairies from two independent biological replicates of H3K27me3, H3 and inputs and three independent biological replicates of H3K9me2, were generated and sequenced on a NextSeq 500 Illumina at the University of Aix-Marseille TAGC U1090 platform. Sequence quality was checked using FASTQC (v0.11.5) (Andrews, 2010). Sequences were trimmed with Trimmomatic (v0.36) (Bolger *et al*, 2014) with the following options: ILLUMINACLIP:TruSeq3-SE.fa with 1 mismatch allowed, and MINLEN:65. Reads were then aligned to the genome using Bowtie v2 (v2.2.9) (Langmead & Salzberg, 2012). The output Sam files were converted to bam with samtools (v1.3.1) (samtools view-bt) using Arabidopsis_thaliana TAIR10.31.dna.genome.fa as a reference. The reads with poor mapping quality were removed with bamtools (v2.4.0) (Barnett *et al*, 2011) and reads with a MAPQ>=20 were kept. The PCR duplicates were removed with the MarkDuplicates tool (v2.8.1) with default options. Peak calling was performed using the diffReps-nb tool (v1.55.2) and bam files were converted to bed files using bedtools (v2.25.0). Peak calling was performed with a window of 200, *p*-value of 0.01 and g test (individual replicate analysis) or negative binomial (analysis with all replicates, Appendix Table S5) statistical tests. The H3 samples of Col-0 and *caa39* were used as background. Annotation was made using PAVIS (Huang *et al*, 2013). For visualization of the data with IGV (v2.3.88) (Robinson *et al*, 2011; Thorvaldsdóttir *et al*, 2013), bigwig files were generated from the alignment files with BamCoverage (v2.4.1) (Ramírez *et al*, 2014). Bam files were indexed with samtools index and then the bigwig files were generated with bamCoverage using the –normalizeTo1x option. The linear K-means clustering method implemented in seqMINER v1.3.4 (Ye *et al*, 2011) was used.

### Extraction of euchromatic island sequences

Those islands harbor characteristic chromatin states of expressed gene and are surrounded by state 9 and 8. Using a homemade perl script, we screened for all regions within heterochromatic regions that are surrounded by chromatin state 8 in the proximal region and state 9 in the distal region, as defined by Sequeira-Mendes et al., 2014. The length threshold of state 9 was set at 450 bp and of the state 8, 150 bp. The list of coordinates generated by this script was then converted to a bed file and used to define the region of interest in the bigwig files. In order to map read density along islands, we used the aggregate command of the bwtool (v1.0) (Pohl & Beato, 2014) with the functionality up:meta:down where meta is set to the median length (in bp) of the element studied.

### Statistical analysis of the Eukaryotic island profiles

The matrix program of Bwtools (<u>h ttps://github.com/CRG-Barcelona/bwtool/wiki/matrix</u>) was used to extract the data from bigwig files. Using the up:meta:down option, the data was extracted in the same way as the aggregate program but instead of returning a mean for each position, it creates a matrix containing each replicate value for each position. The H3K9me2 or H3K27me3 matrix were normalized by the H3 matrix, replicates were concatenated in one matrix which was then treated as the aggregate file. A Shapiro test was first performed to check for normal distribution. The non-parametric Mann-Whitney test was used to compare the value for WT and *caa39* at each position.

### Immunofluorescence

Cytological experiments were conducted as previously described (Bourbousse *et al*, 2015) with a minor modification. For immunolocalization of methylated histones, forty cotyledons were extensively chopped in Galbraith buffer (45 mM MgCl_2_, 20 mM MOPS, 30 mM sodium citrate, 0.3% triton X-100, pH7). Anti-5-methylcytosine (Eurogentec BI-MECY-0100, lot: vt150601) or anti-H3K9me2 (Abcam ab1220, lot: GR244373-6) antibodies were used at 1:100. Anti-mouse antibodies coupled with Alexa 488 (ab150125, lot: GR285477-1) were used at 1:200. At least 15 individual nuclei were observed in each condition and experiments were independently repeated twice.

### Transient transformation and protoplasts preparation

*Agrobacterium*-mediated transient transformation of *Nicotiana benthamiana* was done as previously described (Meteignier *et al*, 2016) with minor modification. Overnight cultures of bacteria were resuspended in infiltration buffer (10mM MgCl_2_, 10mM MES pH5.6, 100µM Acetosyringone) and incubated 2h before infiltration. P19 was added in every infiltration combination. Protoplasts were prepared as previously described (Yoo *et al*, 2007) with minor modifications: enzymes solution was not filtered and protoplasts were not washed.

### Confocal and epifluorescence microscopy

Images were taken on a Zeiss LSM 780 confocal microscope for immunofluorescence experiments or an AxioImager Z1 apotome for every other experiment. All confocal images were acquired using identical parameters.

### Availability of data and material

ChIP-seq, RNA-seq, BS-seq and microarray datasets are available at https://www.ncbi.nlm.nih.gov/geo/query/acc.cgi?acc=GSE103924

## Acknowledgments

We thank Imen Mestiri (IBENS, Paris) for her expertise with cytogenetics. We also want to express our gratitude to students who contributed to this work, especially Justine Quillet, Julien Vieu and César Botella. We thank Ben Field for critical reading of the manuscript. This work was supported by the French National Research Agency (ANR 2010-JCJC-1205-01 and ANR-14-CE02-0010 to CL). Work by FB was supported by the Investissements d’Avenir LabexMemory in Living Systems (MEMOLIFE) grant ANR-10-LABX-54. LD was supported by CEA and Région PACA. High-throughput RNA-sequencing was performed at the POPS plateform, supported by the LabEx Saclay Plant Sciences-SPS (ANR-10-LABX-0040-SPS). MS analysis was performed at the IMM platform supported by a grant from GIS IBiSA. High-throughput ChIP-seq was performed at the TGML platform, supported by grants from Inserm, GIS IBiSA, Aix-Marseille Université, and ANR-10-INBS-0009-10.

## Author contributions

LVM, FV, CV, LD, MR, and CB performed the experiments. CL and MT performed the bioinformatic analyses. LST was in charge of the sequencing. LVM, CL, FB and CL analyzed the data. LVM, KS, FB and CL designed the research. LVM, FB and CL wrote the manuscript. All authors read and approved the final manuscript.

