## Supplemental figures for "Topoisomerase VI participates in an insulator-like function that prevents H3K9me2 spreading into euchromatic islands"

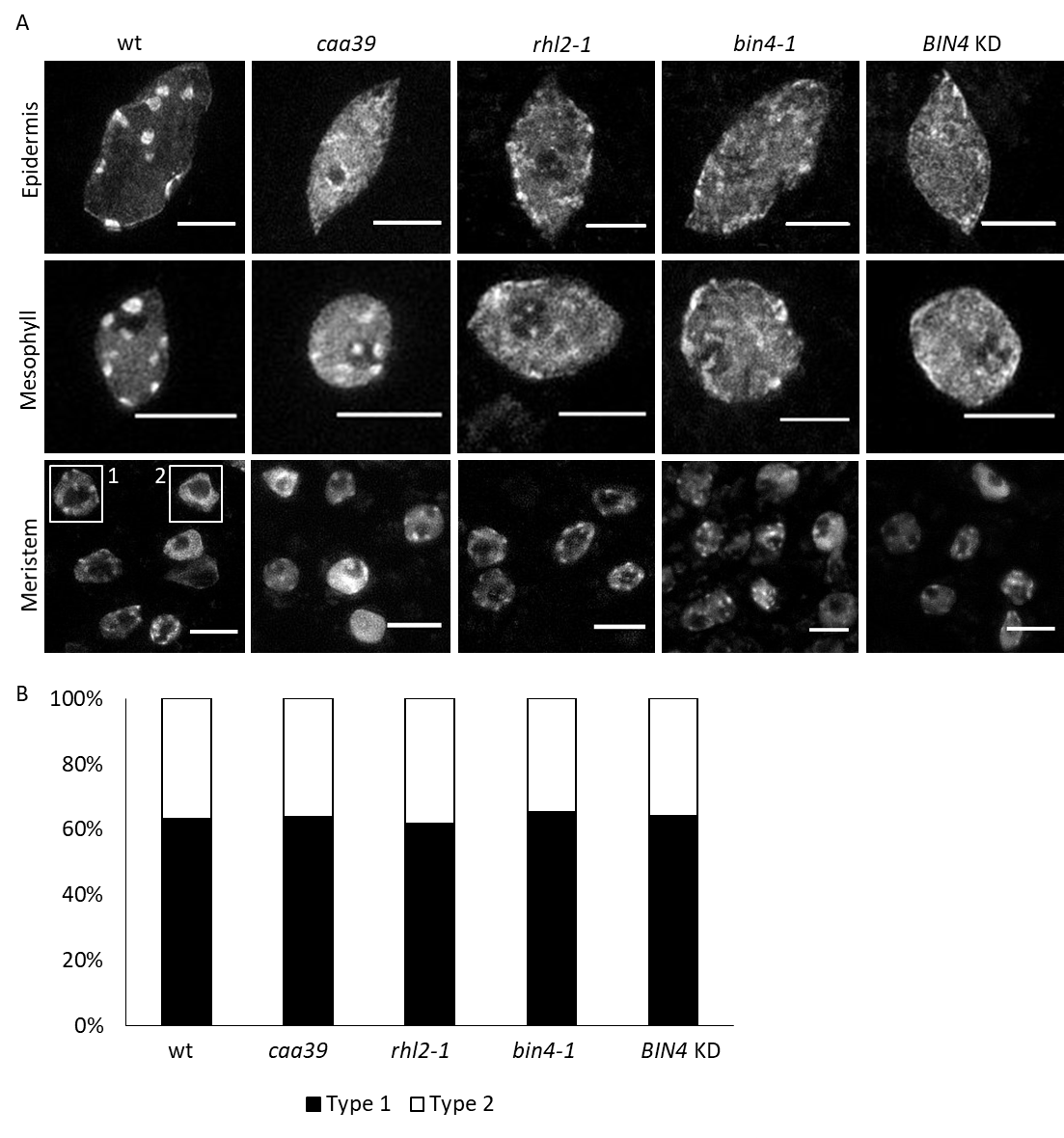
**Figure S1. Nuclear phenotype of Topo VI mutants in post-mitotic and mitotic cells of aerial parts.** (A) Representative Z-projections of apotome micrographs of DAPI-stained nuclei from fixed tissues of the indicated genotypes and cell types. Scale: 10 µm. (B) Proportion of nuclei harbouring condensed chromocenters (type 1) or decondensed chromocenters (type 2) in nuclei of mitotic cells of shoot apical meristems of indicated genotypes. Number of nuclei in wt: 38, *bin4-1*: 49, *rhl2-1*: 34, *caa39*: 36, *BIN4* KD: 75

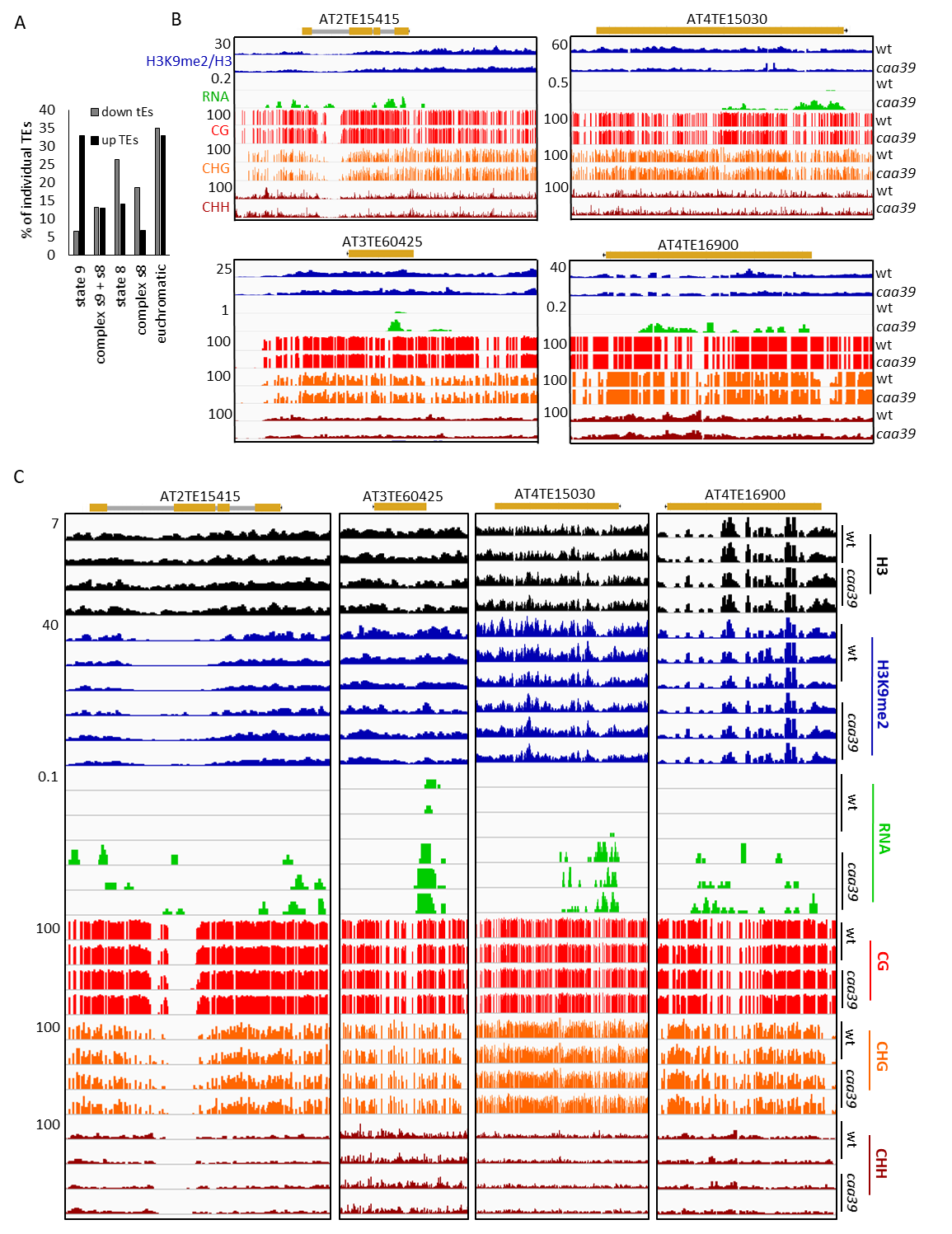

**Figure S2. Topo VI is required for the silencing of heterochromatic (state 9) transposable elements.** (A) Chromatin state features of down- and up-regulated TEs. (B) IGV screenshots of representative heterochromatic TEs. (C) IGV screenshots of representative heterochromatic TEs from individual experimental replicates.

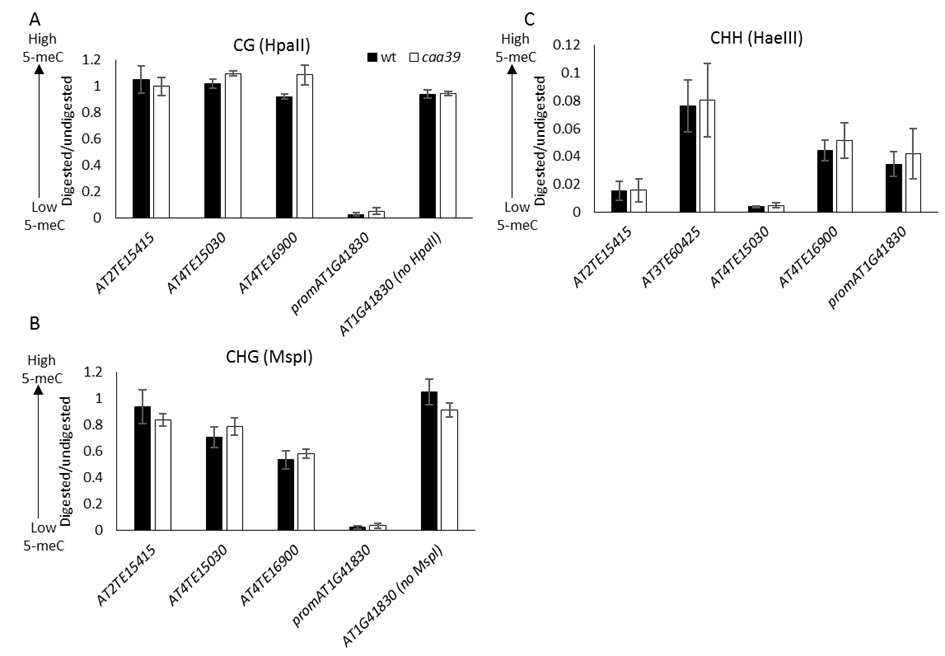

**Figure S3. Analysis of DNA methylation in wt and the Topo VI hypomorphic mutant *caa39*.** Genomic DNAs were extracted and submitted or not to HpaII (a), MspI (b) or HaeIII (c) digestion. The promoter region of the pericentromeric gene *AT1G41830* was used as a poorly methylated control locus, and primers spanning no HpaII/MspI restriction site in that gene were used as a negative control. Error bars: ±SEM of three biological replicates. Note there is no MspI/HpaII restriction site in *AT3TE60425*.

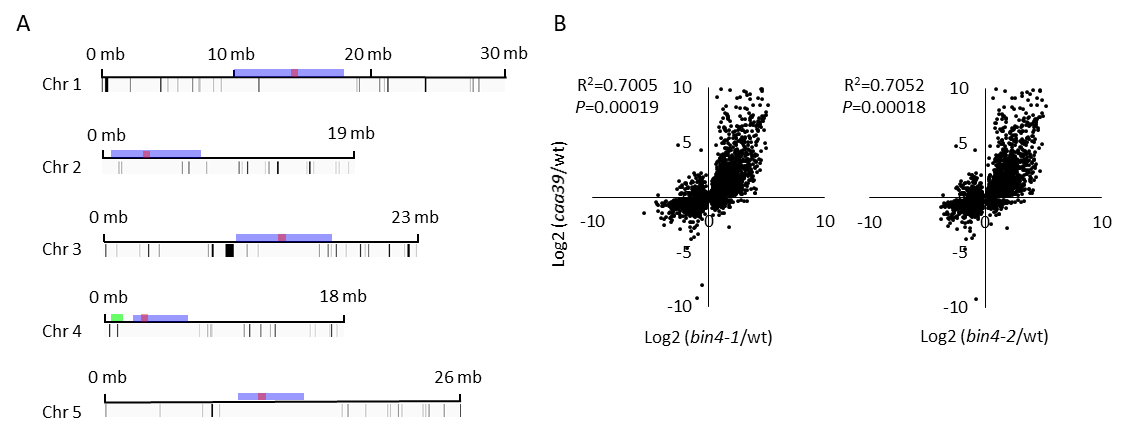

**Figure S4. Gene expression changes in Topo VI mutants in relation with H3K9me2 levels**. (A) Positional gene enrichment analysis of the top 500 upregulated genes in *caa39*. Black lines correspond to enriched regions, blue boxes correspond to pericentromeric regions as defined by Yelina et al. (2012), green box to the knob and red to centromeres. (B) Scatter plots and pearson correlations of differentially expressed (*P* < 0.05) genes in *caa39* and *bin4-1* or *bin4-2*.

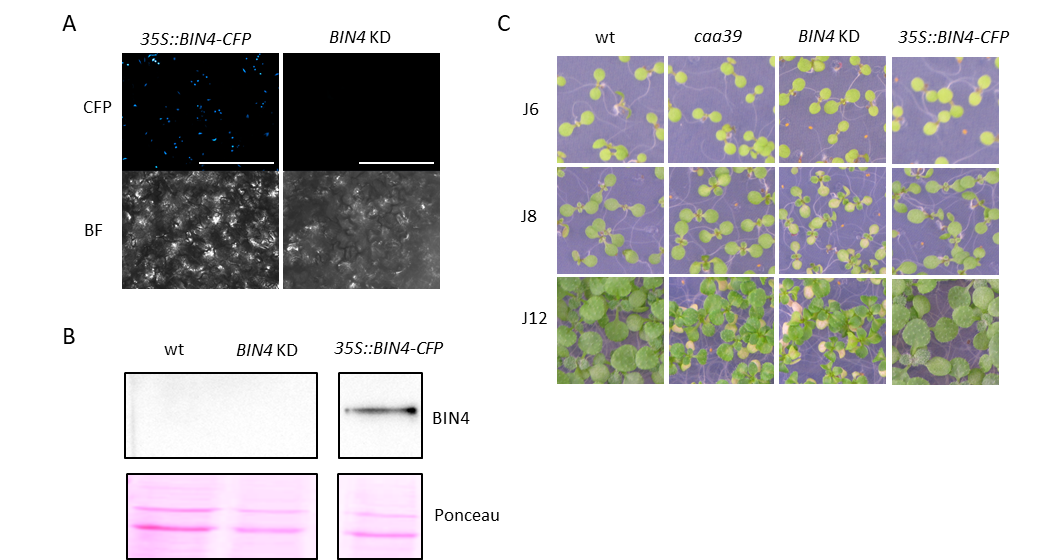
 **Figure S5. A *BIN4* co-suppressed transgenic line (*BIN4* KD) develops a phenotype similar to *caa39*** (A) Representative nuclear distribution of BIN4-CFP in cotyledons of six day-old *35S::BIN4-CFP* transgenic plants. Fluorescence is absent in the KD line. (B) Western blot analysis of the accumulation of BIN4-CFP fusion protein in KD lines and *35S::BIN4-CFP* transgenic plants. Ponceau staining is shown as a loading control. (C) Representative phenotypes of *in vitro*-grown wt, *caa39*, *BIN4* KD and *BIN4-CFP* overexpressing lines 6, 8, and 12 days post-germination.

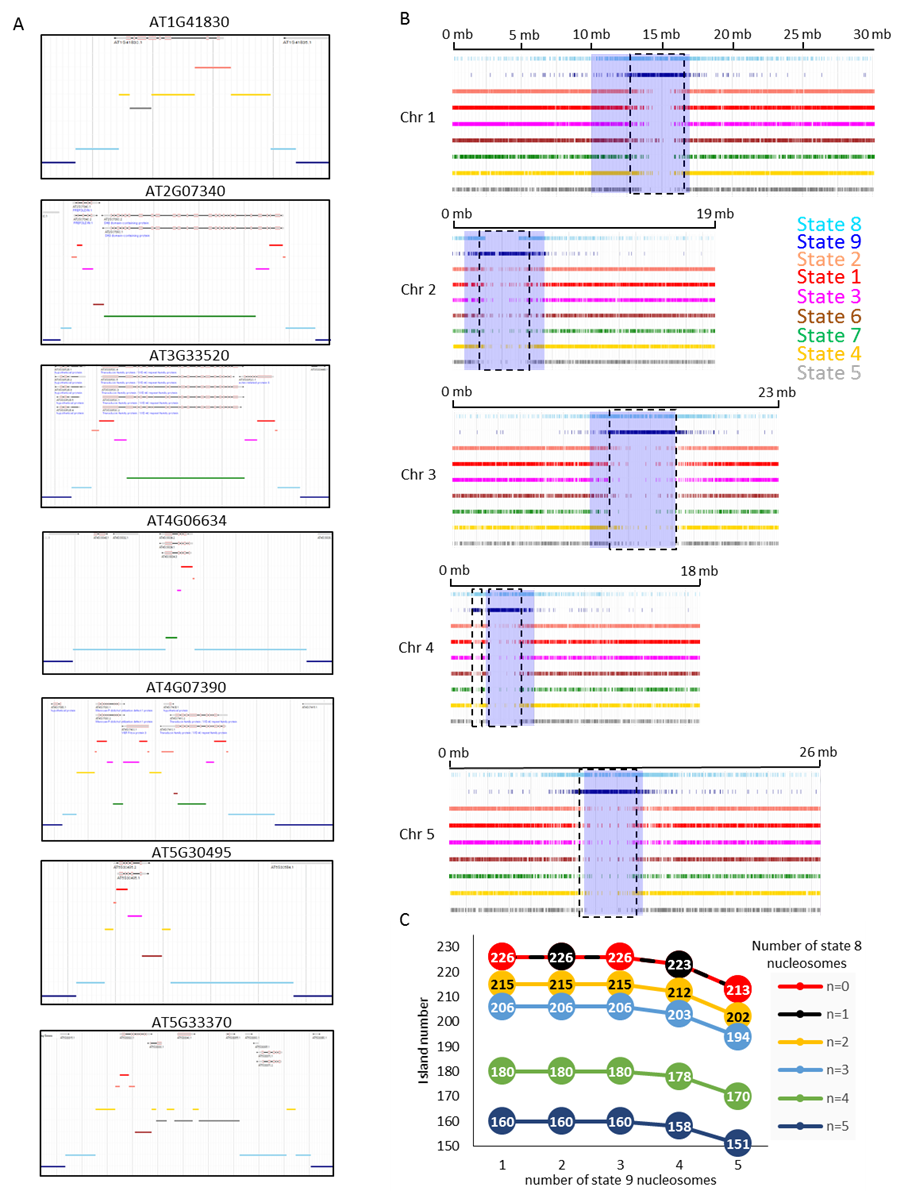
 **Figure S6.**  **Features of euchromatic islands.** (A) Screenshots of the Araport JBrowse tool showing the chromatin context of seven representative EI genes validated by RT-qPCR (B) Araport JBrowse screenchots of chromatin states at the level of chromosomes. The black dashed frame indicates heterochromatic regions and the light purple boxes indicate pericentric regions as defined by Yelina *et al.* (2012). Coordinates of dashed frames; chr1: 13 230 300-16 753 200; chr2: 2 362 500-5 636 850; chr3: 11 313 900-15 740 700; chr4 knob: 1 591 800-2 367 300; chr4: 2 782 500-5 142 900; chr5: 10 348 800-13 583 850. (C) Number of extracted EIs relative to the number of nucleosomes in chromatin states 8 and 9 specified in the extraction script. Each curve corresponds to a different number of state 8 nucleosomes.

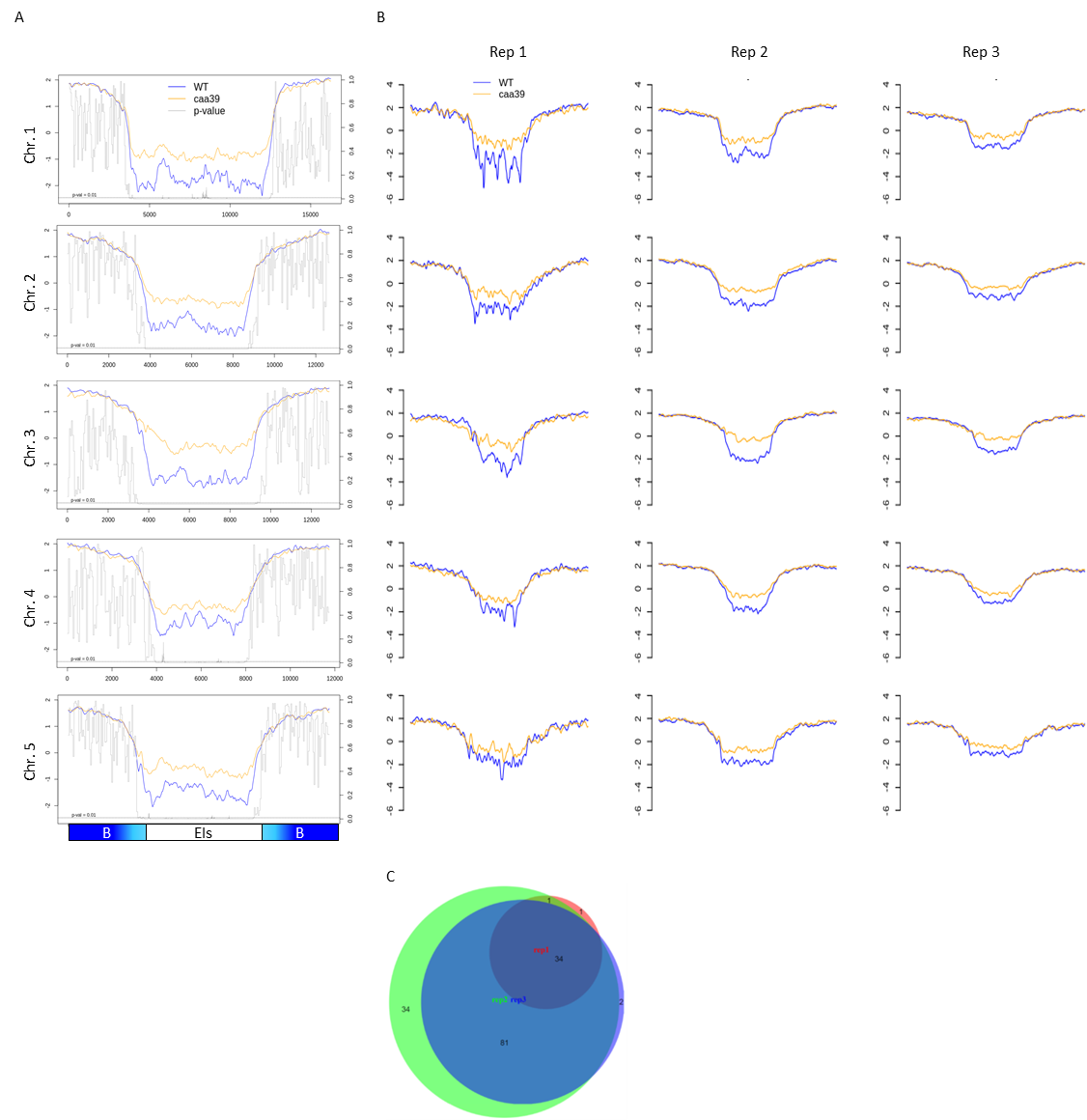

**Figure S7. Average distribution of H3-normalized H3K9me2 along euchromatic gene islands and 4 kb flanking regions in wt and *caa39*.** (A) Left Y axis represents average Log2(H3K9me2/H3) along euchromatic gene islands and 4 kb in surrounding borders for each chromosome. Two (H3) and three (H3K9me2) independent biological replicates for each genotype were performed. The right y axis represents the *P*-value. *P*-value was calculated according to the Mann-Whitney test. (B) Log2(H3K9me2/H3) is shown for each independent biological replicate and each chromosome. (C) Venn diagram of common EIs between replicates. EIs containing differential H3K9me2 peaks in *caa39* versus wt were identified using diffReps in individual replicates and compared between replicates.

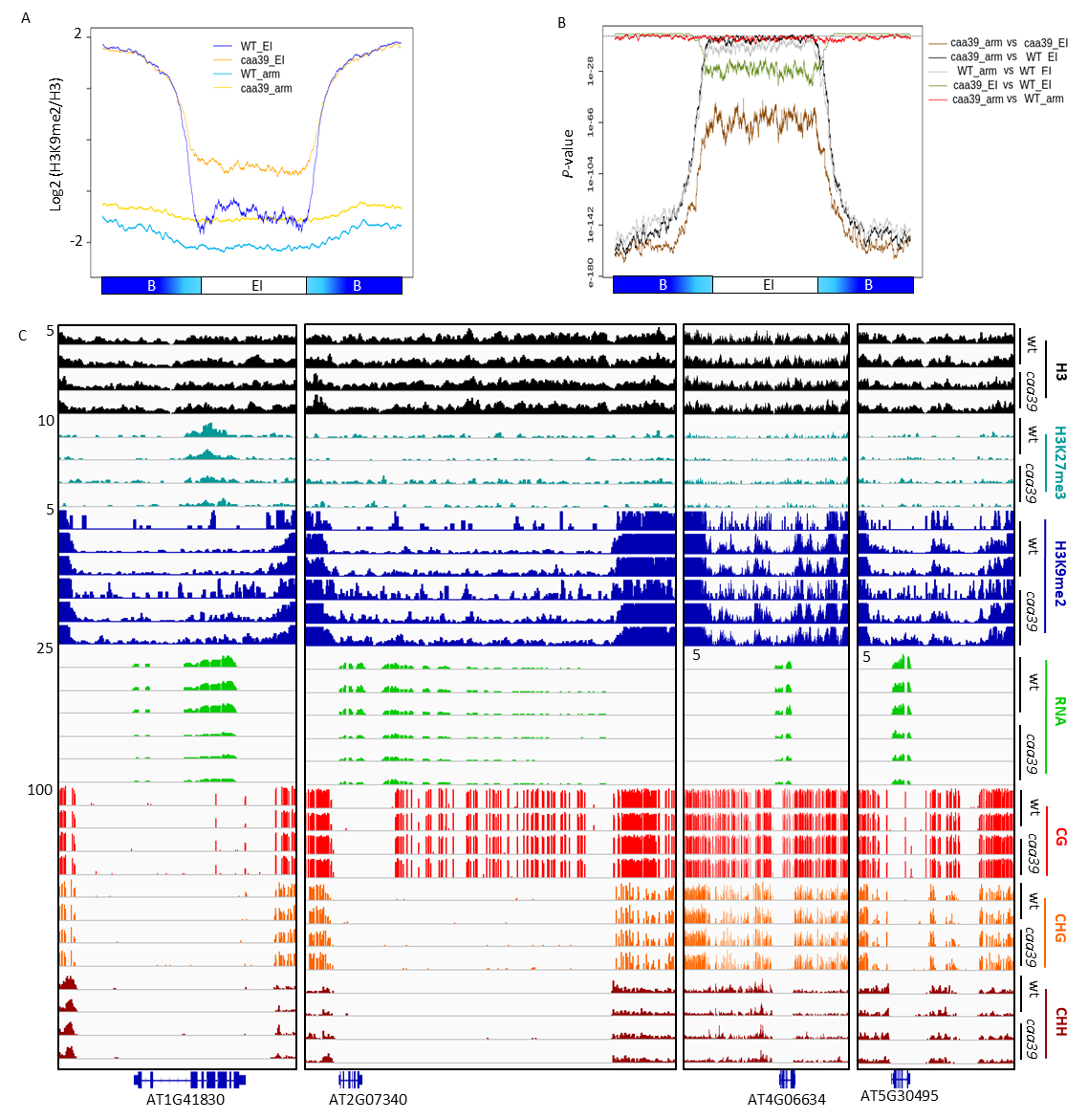

**Figure S8. Topo VI inhibits H3K9me2 spreading predominantly in pericentromeric regions.** (A) Average distribution of H3-normalized H3K9me2 along state 8-free EIs and random euchromatin regions of chromosome arms, and 4 kb flanking regions. Two (H3) and three (H3K9me2) independent biological replicates for each genotype were performed. Light blue and light orange represent the ±SEM. (B) *P*-value was computed for each aggregated point in (A) by using a Mann-Whitney test. (C) IGV screenshots of four individual small s8-free EIs from individual experimental replicates.

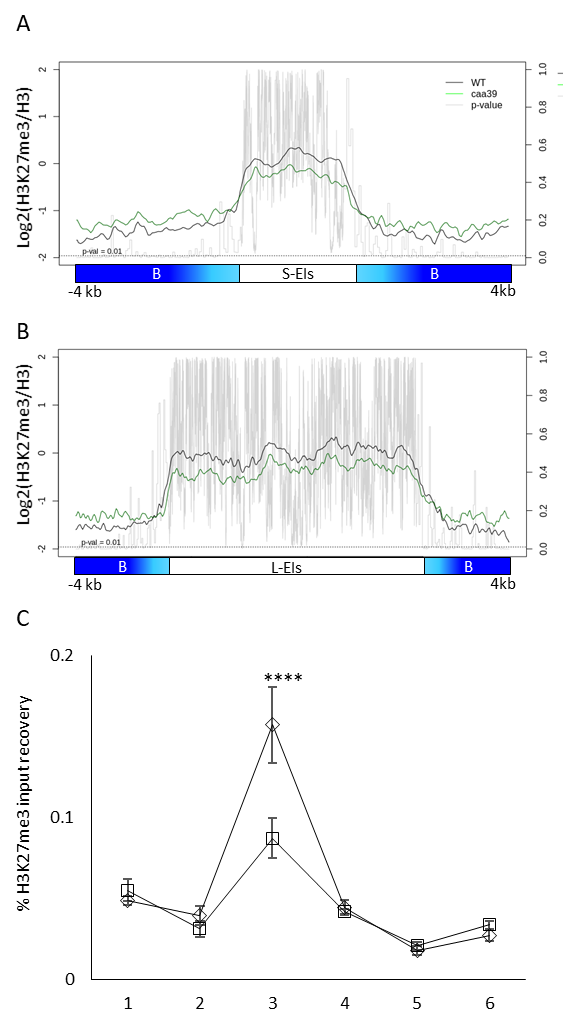

**Figure S9. Average distribution of H3-normalized H3K27me3 along euchromatic gene islands in wt and *caa39*.** (A) Log2(H3K27me3/H3) in small or large (B) euchromatic islands. The right y axis represents the *P*-value. *P*-value was calculated according to the Mann-Whitney test. (C) ChIP-qPCR validation of H3K27me3 decrease in a single gene-containing island. Error bars: ±SEM of three biological replicates. ****: P < 0.0001 (two-way ANOVA, Fisher’s test).

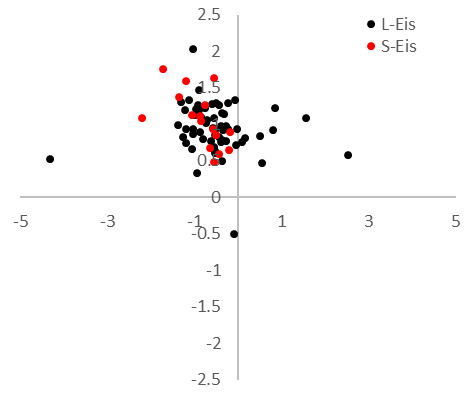

**Figure S10. Scatter plot of RNA-seq Log2(FC) and Log2(H3K9me2/H3).**

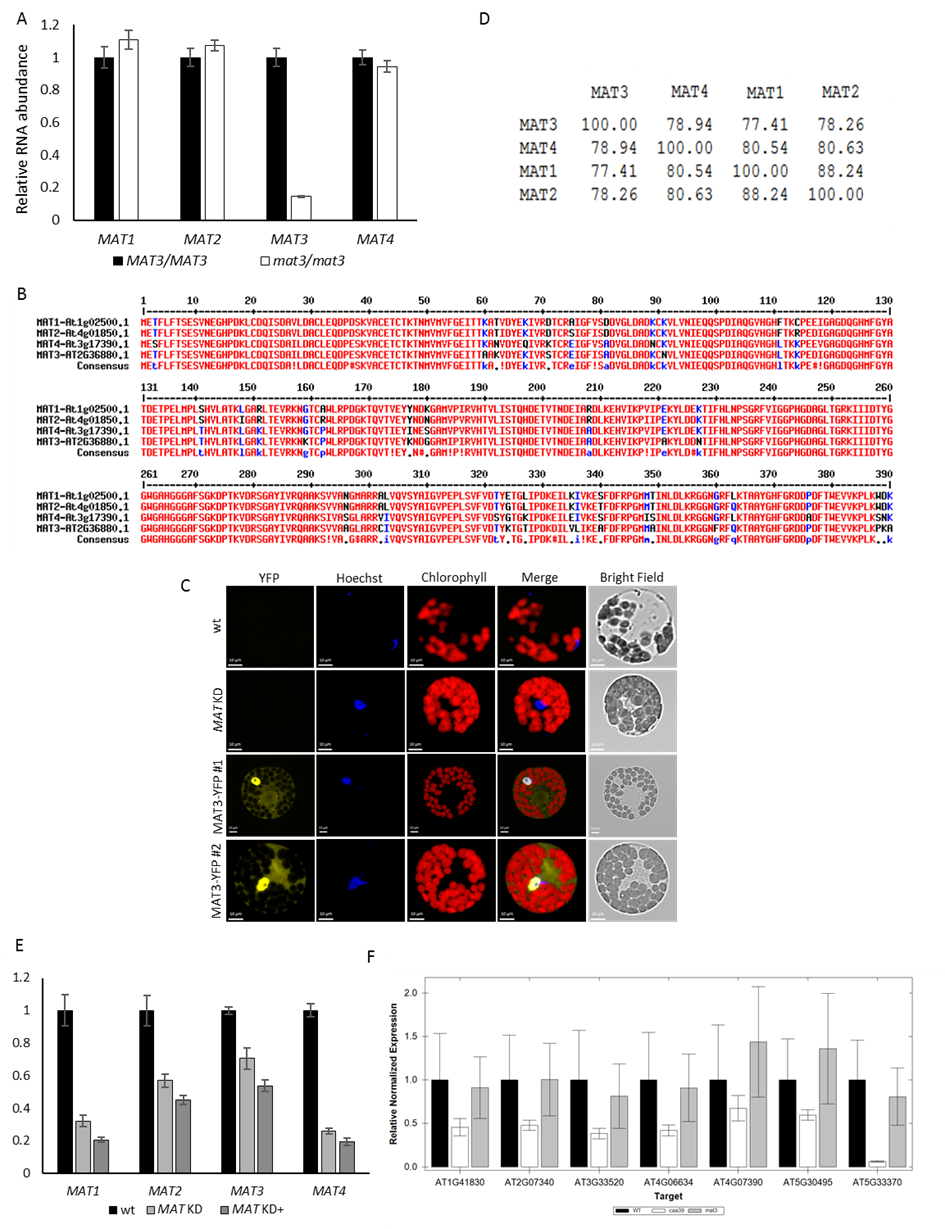

**Figure S11. Characterization of the *mat3* mutant and *MATs* KD lines.** (A) RT-qPCR experiment showing the amount of *MAT* transcripts in the *mat3* mutant compared to a WT isogenic line. Error bars: ±SEM of two biological replicates. (B) Amino acid sequence alignment of MAT isoforms in *Arabidopsis*. (C) Protoplasts from stable overexpressing transgenic lines were prepared, stained with Hoechst 33342 and imaged for YFP, Hoechst and bright field. The localization of the YFP-MAT3 protein is revealed by merging the Hoechst and YFP channels. (D) Percent of MATs CDS sequences identity in *Arabidopsis*. (E) RT-qPCR analysis of *MAT1/2/3/4* transcript abundance in *MATs* KD line and wt. Error bars: ±SEM of three biological replicates. (F) RT-qPCR analysis of the seven EI genes in indicated genotypes. Error bars: ± SEM of three biological replicates.

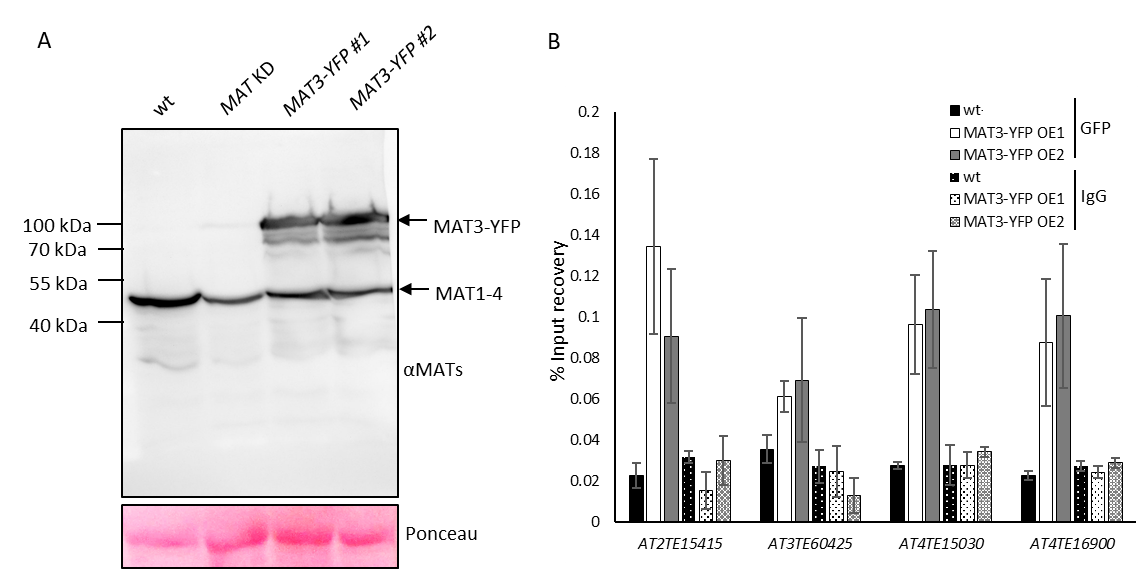

**Figure S12.** **Validation of anti-MAT antibody and TEs enrichment in MAT3-YFP ChIPs.** (A) Total protein extracts of the indicated genotypes were separated on SDS-PAGE gel, transferred to PVDF, and blotted with MAT1-4 antibody. Ponceau staining is shown as a loading control. (B) Chromatin of 6 day-old wt or MAT3-YFP stable transgenic cotyledon nuclei was IPed with an anti-GFP antibody and the recovery of TEs awakened in *MAT* KD and *caa39* was measured by qPCR. Error bars: ±SEM of three biological replicates.
