## Appendix Table S4 for "Topoisomerase VI participates in an insulator-like function that prevents H3K9me2 spreading into euchromatic islands"

Total EIs

|  | chr1 | chr2 | chr3 | chr4 | chr5 | All chr |
| --- | --- | --- | --- | --- | --- | --- |
| moyenne | 13146 | 5636 | 8476 | 7158 | 7506 | 8494.28 |
| médiane | 8249 | 4649 | 4874 | 3749 | 5549 | 5099 |
| meta value | 8250 | 4650 | 4880 | 3750 | 5550 | 5100 |

Large EIs

|  | chr1 | chr2 | chr3 | chr4 | chr5 | All chr |
| --- | --- | --- | --- | --- | --- | --- |
| moyenne | 18032.87 | 9101.63 | 16252.57 | 14763.29 | 11767.18 | 14496.87 |
| médiane | 13049 | 7799 | 10724 | 11099 | 10124 | 10724 |
| méta value | 13050 | 7800 | 10725 | 11100 | 10125 | 10725 |

Small EIs

|  | chr1 | chr2 | chr3 | chr4 | chr5 | All chr |
| --- | --- | --- | --- | --- | --- | --- |
| moyenne | 3049 | 2892.75 | 2524.66 | 2337.64 | 2572.68 | 2640.02 |
| médiane | 2699 | 3074 | 2099 | 2099 | 3149 | 2549 |
| méta value | 2700 | 3075 | 2100 | 2100 | 3150 | 2550 |

S8-containing EIs

|  | chr1 | chr2 | chr3 | chr4 | chr5 | All chr |
| --- | --- | --- | --- | --- | --- | --- |
| mean | 16084 | 8804 | 18019 | 16364 | 12664 | 13896 |
| median | 13349 | 7274 | 13799 | 12074 | 10649 | 11699 |
| Meta aggregate value | 13350 | 7280 | 13800 | 12080 | 10650 | 11700 |

S8-free EIs

|  | chr1 | chr2 | chr3 | chr4 | chr5 | All chr |
| --- | --- | --- | --- | --- | --- | --- |
| mean | 10679 | 4676 | 3704 | 3749 | 4205 | 5705 |
| median | 5549 | 3899 | 2549 | 2399 | 3449 | 3524 |
| Meta aggregate value | 5550 | 3900 | 2550 | 2400 | 3450 | 3525 |
